# Estimating movement rates between Eurasian and North American birds that are vectors of avian influenza (AI)

**DOI:** 10.1101/2021.09.10.459847

**Authors:** Fern Spaulding, Jessica F. McLaughlin, Travis C. Glenn, Kevin Winker

**Affiliations:** University of Alaska Museum, University of Alaska Fairbanks, Fairbanks, AK 99775, USA; Department of Biology and Wildlife, University of Alaska Fairbanks, AK 99775, USA; Sam Noble Oklahoma Museum of Natural History, University of Oklahoma, Norman, OK 73072, USA; Department of Environmental Health Science, University of Georgia, Athens, GA 30602, USA

**Keywords:** avian influenza, Beringia, waterfowl, gene flow, subsistence harvest, vector species, Alaska

## Abstract

Avian influenza (AI) is an emerging zoonotic disease that will likely be involved in future pandemics. Because waterbird movements are difficult to quantify, determining the host-specific risk of Eurasian-origin AI movements into North America is challenging. We estimated relative rates of movements, based on long-term evolutionary averages of gene flow, between Eurasian and North American waterbird populations to obtain bidirectional baseline rates of the intercontinental movements of these AI hosts. We used population genomics and coalescent-based demographic models to obtain these gene-flow-based movement estimates. Inferred rates of movement between these populations varies greatly among species. Within dabbling ducks, gene flow, relative to effective population size, varies from ∼3-24 individuals/generation between Eurasian and American wigeons (*Mareca penelope – M. americana*) to ∼100-300 individuals/generation between continental populations of northern pintails (*Anas acuta*). These are evolutionary long-term averages and provide a solid foundation for understanding the relative risks of each of these host species in potential intercontinental AI movements. We scale these values to census size for evaluation in that context. In addition to being AI hosts, many of these species are also important in the subsistence diets of Alaskans, increasing the risk of direct bird-to-human exposure to Eurasian-origin AI virus. We contrast species-specific rates of intercontinental movements with the importance of each species in Alaskan diets to understand the relative risk of these taxa to humans. Greater scaup (*Aythya marila*), mallard (*Anas platyrhynchos*), and northern pintail (*Anas acuta*) were the top three species presenting the highest risks for intercontinental AI movement both within the natural system and through exposure to subsistence hunters. These directly comparable, species-based intercontinental movement rates and relative risk rankings should help in modeling, monitoring, and mitigating the impacts of intercontinental host and AI movements.

## 1.2 Introduction

Avian influenza (AI) is an emerging zoonotic disease and will likely be involved in future pandemics (Webster et al., 1992; Ferguson et al., 2006; Cooper et al., 2006; Sellwood et al., 2007). AI research and surveillance has demonstrated the exchange of viruses between Eurasia and North America through migratory birds that occur across the Beringia region of the North Pacific Ocean (Pearce et al., 2009; Ramey et al., 2010, 2018; Reeves et al., 2013, 2018). Beringia (Fig. 1) encompasses a large geographic area, and many birds migrate seasonally here to breed. Aquatic birds, such as waterfowl and shorebirds, are often asymptomatic carriers of the disease, indicating that they are well adapted to this pathogen and serve as a natural reservoir (Horimoto & Kawaoka, 2001). Winker & Gibson (2010) estimated that ∼1.5-2.3 million birds that are likely vector species of AI migrate from Eurasia across Beringia to their breeding grounds in Alaska every year. This large overlap of Eurasian and North American migration systems causes extensive seasonal contact between these lineages and populations, resulting in a potential for virus movements into and out of North America (Winker et al., 2007; Flint et al., 2009). This vast number of seasonal migrants is often underappreciated when modeling AI introduction into North America (Hinshaw et al., 1980, Donis et al., 1989; Webster et al., 1992; Böning-Gaese et al., 1998; Kilpatrick et al., 2006; Rappole et al., 2006; Winker et al., 2007; Winker & Gibson, 2010). International borders and vast areas of remote land make accurate estimates of intercontinental AI host movements difficult. Yet knowing the degree of these movements between Eurasia and North America is critical to understanding intercontinental transmission of AI viruses.

**Figure 1.**
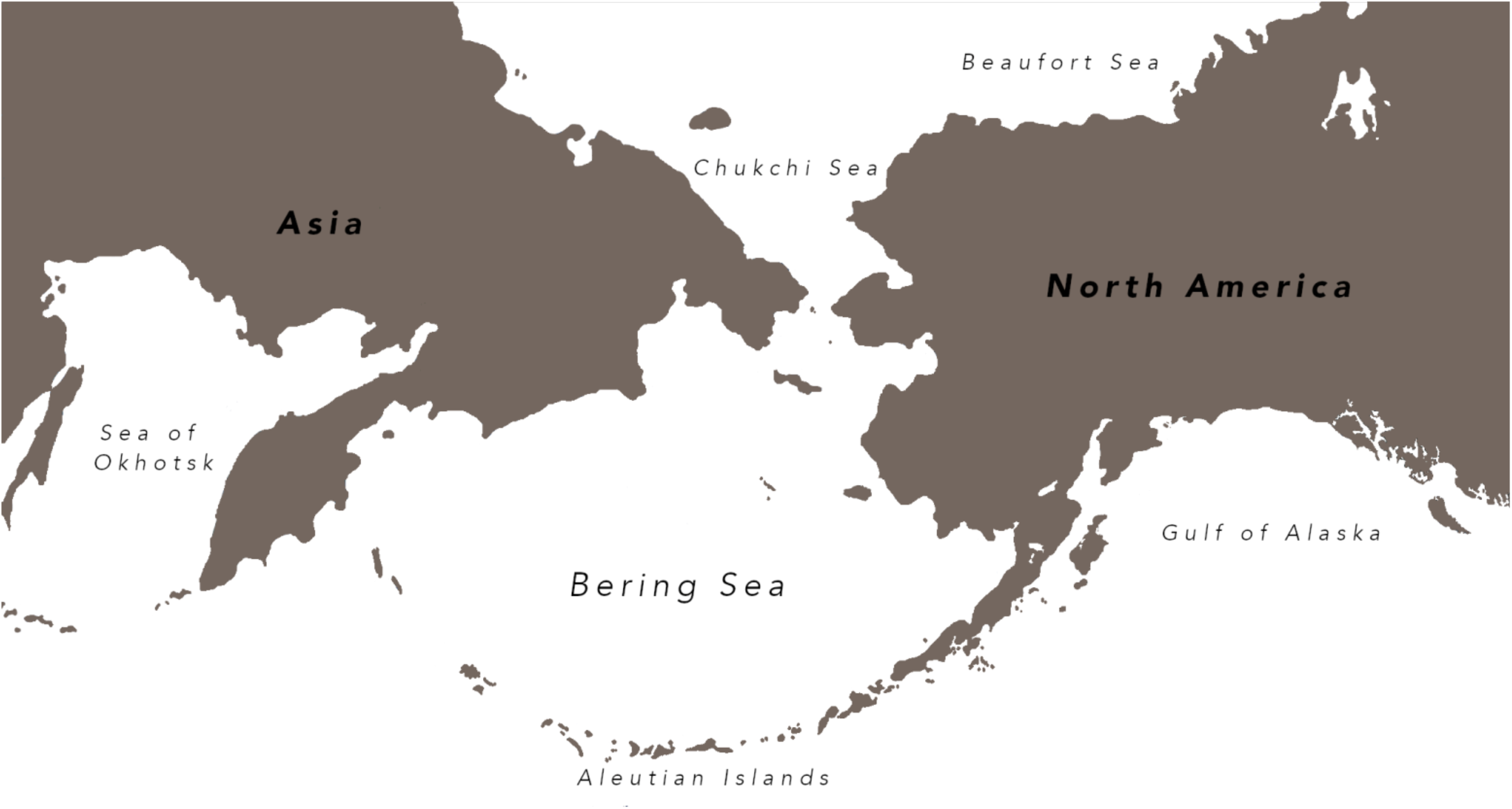
Beringia encompasses the region extending from the Russian Far East across the Bering Sea though Alaska to western Canada in North America. Across Beringia, waterbirds are important avian influenza (AI) vectors and are a staple in the rural subsistence diet.

In addition to being potential vectors of intercontinental virus transport, many of these waterfowl and shorebird species are a common staple in the subsistence diets of many Alaskans. Waterfowl in particular account for a large proportion of annual harvest, making up approximately 85% of migratory birds taken for subsistence in Alaska (Naves & Keating, 2017). Many of these birds are possible carriers of AI, including the highly pathogenic avian influenza (HPAI) of Eurasian-origin that has been detected in Alaskan populations of waterfowl (e.g. H9N2, Ramey et al., 2015), and AI viral dispersal across Beringia is not a rare occurrence (Ramey et al., 2018). Extensive human contact with wild populations of waterfowl during hunting might cause AI exposure, a possibility supported by the detection of antibodies in humans to influenza virus subtypes (i.e., H11) found only in wild birds (Gill et al., 2006). Hunters will often process harvested birds themselves while in the field without using gloves or personal protective equipment (Dishman et al., 2010). By processing an influenza-infected duck, a hunter might be exposed to virus-laden mucosa and excretions (i.e., nasal or fecal) in addition to blood, tissues, and other body fluids (Brown et al., 2007; Siembieda et al. 2008; Dishman et al., 2010). Because waterfowl are an important subsistence food for Alaskans, contact while handling and processing these birds is a potential risk factor for direct bird-to-human transmission (Gilsdorf et al., 2006; Yamamoto et al., 2010; Dórea et al., 2013).

In this study, we used population genomics, using ultraconserved elements (UCEs) as a sequenced subsampling of the genome, to improve rough estimates (Winker & Gibson, 2010) of intercontinental AI host movements among key waterbird species to better understand the natural host-virus, movement-transmission landscape in Beringia. Given Alaska’s proximity to Eurasia, we also contrast these estimates with the importance of each species in Alaskan diets to understand the relative risk of these taxa directly to human consumers. This information can enable subsistence hunters to selectively harvest species that have a lower potential risk of carrying Eurasian-origin HPAI. These relative risk rankings should help in modeling, monitoring, and mitigating the impacts of intercontinental host and AI movements.

## 1.3 Materials and Methods

### 1.3.1 Sampling and laboratory

Our samples consist of high-quality, vouchered tissue samples from wild birds of Eurasian and North American origin (Table S1). Our study design includes pairwise comparisons of populations, subspecies, and species, because taxonomy is not a reliable indicator of intercontinental levels of gene flow (Humphries & Winker, 2011; Peters et al., 2012, McLaughlin et al., 2020). We sampled the following numbers of individuals for each vector species: 10 northern pintail (*Anas acuta*), 12 mallard (*Anas platyrhynchos*), 10 greater scaup (*Aythya marila*), 10 common eider (*Somateria mollissima*), and 9 common and Wilson’s snipe (*Gallinago gallinago – G. delicata*). We also incorporated the gene flow rates obtained by McLaughlin et al. (2020) which include: green-winged teal (*Anas crecca crecca – A. c. carolinensis*), long-tailed duck (*Clangula hyemalis*), and Eurasian and American wigeons (*Mareca penelope* -*M. americana*). Using UCEs as a sequence-based subsampling of the genome allows us to examine thousands of orthologous loci (Faircloth et al., 2012), and it has been shown that, in general, relatively small sample sizes are sufficient when using coalescent methods to estimating population demographics (Felsenstein, 2005; Nazareno et al., 2017; McLaughlin & Winker, 2020). Here, we are particularly interested in levels of gene flow (*m*), which when using our methods of analysis appear to be relatively consistent even when sample sizes vary (McLaughlin & Winker, 2020). Therefore, we consider our sample sizes to be adequate for the questions we pose. DNA extractions followed standard protocol for animal tissues using the QIAGEN DNeasy Blood + Tissue Extraction Kit (QIAGEN, 2006). Dual-indexed DNA libraries were prepared and quantified using Qubit fluorimeter (Invitrogen, Inc., Carlsbad, CA, USA). The samples were enriched for 5,060 ultraconserved element (UCE) loci using the Tetrapods-UCE-5Kv1 kit from MYcroarray following version 1.5 of the UCE enrichment protocol and version 2.4 of the post-enrichment amplification protocol (http://ultraconserved.org/). The resulting pool was sequenced using a paired-end 150 (PE150) protocol on an Illumina HiSeq 2500 using three lanes (Illumina, Inc., San Diego, CA, USA; UCLA Neuroscience Genomics Core).

### 1.3.2 Bioinformatics

Raw and untrimmed FASTQ sequence data that contained low-quality bases were removed using Illumiprocessor (v.2.0.6; Faircloth, 2013). We built UCE reference sequences for each vector species by combining sequence read files (read1 plus singletons and read2) from two individuals (resulting in one read 1 and one read 2 file). We used the program Trinity (v.2.4.0; Grabherr et al., 2011) to assemble these reads *de novo* on the Galaxy platform (v.2.4; Afgan et al., 2016), found and extracted UCE loci from this assembly using PHYLUCE (v.1.5.0; Faircloth, 2016) by matching the contigs to the probes set used, and then saved the resulting sequences as a reference FASTA. SNPs within the reference were coded using IUPAC codes. Our bioinformatics pipeline focused on the package PHYLUCE, which calls many dependencies and identifies conserved orthologous loci that are then used as our reference set of UCE loci to call individual variants. Briefly, individual read1 (plus singletons) and read2 files were mapped to the taxon-specific reference and indexed using BWA-MEM (v.0.7.7; Li & Durbin, 2009; Li, 2013) and SAMtools (v.0.1.19; Li et al., 2009). Next, PICARD (v.1.106; http://broadinstitute.github.io/picard) was used to clean the alignments, add read group header information, and remove PCR and sequencing duplicates. SNPs were called for each individual against the reference sequence using Genome Analysis Toolkit module UnifiedGenotyper (GATK, v.3.3.0; McKenna et al., 2010). GATK was also used to call and realign around indels, call and annotate SNPs, filter SNPs around indels, and then restrict data to high-quality SNPs. VCFtools (v.0.1.13; Danecek et al., 2011) was used to filter the high-quality SNPs to create a complete matrix (all individuals represented at all loci) with a minimum genotype quality (Phred) score of 10.0 (which equates to 90% confidence). The high-quality VCF was then thinned to 1 SNP/locus and made biallelic. We used BLASTn on NCBI to identify sex-linked loci (Z-linked) using our high-quality reference data of confidently surveyed loci, and resulting hits for Z-linked loci were then removed using a custom script (find_chromy v.1.2; https://github.com/jfmclaughlin92/beringia_scripts/). Our thinned, biallelic VCF with Z-linked loci removed was used for our demographic analyses to estimate movement (i.e., gene flow).

### 1.3.3 Demographic analyses to estimate gene flow

Diffusion Approximations for Demographic Inference (δaδi, v.1.7.0; Gutenkunst et al., 2009) was used to estimate demographic parameters under best-fit models of pairwise divergence. Note that in population genomics models, “migration” equals gene flow and is not related to the seasonal migration of individuals. To find a best-fit model of demographic history, we tested eight divergence models (Fig. S1): A) no divergence (neutral, populations never diverge); B) split with no migration (divergence without gene flow); C) split with migration (divergence with gene flow that is bidirectionally symmetric, 1 migration parameter); D) split with bidirectional migration (divergence with gene flow that is bidirectionally asymmetric, 2 migration parameters); E) split with exponential population growth, no migration; F) split with exponential population growth and migration; G) secondary contact with migration (1 migration parameter); and H) secondary contact with bidirectional migration (2 migration parameters). The scripts for these models are available here: https://figshare.com/s/e75158104c7896fa79e7.

For each pairwise comparison, we ran a series of optimization runs that consisted of running the model repeatedly to fine-tune model parameters. Following optimization, the best five log-likelihood scores from each set of subsequent runs were averaged to summarize that model. We used the Akaike Information Criterion (AIC; Akaike, 1974; Burnham & Anderson, 2004) to determine the best-fit model. Once the best-fit model was determined, it was run at least 15 times to obtain demographic parameter estimates, and we used these estimates from the best-fit model’s top three runs to calculate biological estimates. Then the model was bootstrapped to provide a 95% confidence interval around each demographic parameter. To convert the best-fit model’s demographic parameters to biologically relevant values, we determined generation time from the literature and estimated substitution rates (Table S2) using BLASTn to compare each reference FASTA to a related avian genome with a fossil-calibrated node (Claramunt & Cracraft, 2015). These values were then used with the best-fit model demographic parameter estimates obtained from our analyses to provide estimates of ancestral population size (*N*_*ref*_), size of populations (*nu1, nu2*), time since split (*T*), migration (*M*; gene flow in individuals/generation, derived from the raw model output *m*), migration from population 1 into population 2 (derived from *m12*), migration from population 2 into population 1 (derived from *m21*), and time of secondary contact (*T*_*sc*_) as appropriate (based on the best-fit model for each pairwise comparison).

### 1.3.4 Scaling movement, harvest rates, and risk factors

Estimates of gene flow from the demographic analysis in δaδi are based on long-term effective population sizes (*N*_*e*_), which is nearly always lower than census size in high-latitude birds (e.g., McLaughlin et al., 2020). These evolutionary average rates of gene flow are not scaled to census size, so they are very conservative, but they are directly comparable to each other. To consider these rates relative to census sizes, estimates of the latter were obtained from population estimates from the U.S. Fish and Wildlife Service (USFWS, 2019) and from Wetlands International (2021). The USFWS combines both greater and lesser scaup (*Aythya marila* and *Aythya affinis)* into one census estimate because the two species can be difficult to distinguish. To address this, we used the ratio of greater and lesser scaup obtained from the Bird Conservancy of the Rockies’ global population estimates as part of their Partners in Flight, Avian Conservation Assessment Database (ACAD, 2020). We applied this ratio to the USFWS (2019) census estimate of “scaup” to obtain a census estimate for greater scaup alone. We used the proportion of gene flow (*M*, in individuals/generation) relative to effective population size (*N*_*e*_) to scale our estimates of long-term gene flow up to approximately estimate contemporary gene flow by multiplying this proportion by census size: (*M* _*Eurasia*_/*N*_*e North America*_*\**census size).

Subsistence harvest rates in Alaska were obtained from Naves & Otis (2017) and Naves & Keating (2018, 2020). Some subsistence species were not categorized with a species name, so the species categorized as “Teal” were used for green-winged teal (*Anas crecca crecca – A. c. carolinensis*), “Scaup” for greater scaup (*Aythya marila*), and “Small Shorebird” for common and Wilson’s snipe (*Gallinago gallinago – G. delicata*). We calculated the average annual harvest for each vector species across the years 2016, 2017, and 2018. To calculate a relative risk of potential exposure of Eurasian-origin AI virus to subsistence users, we estimated the number of individuals harvested of Eurasian origin given (*M/N*_*e*_) and annual harvest (*M* _*Eurasia*_*/N*_*e*_*\**annual harvest). Given high variation in AI infection rates (e.g., annual, seasonal, geographic), we did not add this as a variable.

## 1.4 Results

### 1.4.1 Estimated rates of gene flow

The best-fit models for our demographic analyses in δaδi found gene flow present in all pairwise comparisons (Table S3). In half of our comparisons (four species) the AIC analyses showed that some models fit the data similarly well, yielding a best-fit model and a runner-up model that was not statistically worse. These cases were in green-winged teal (*Anas crecca crecca* – *A. c. carolinensis*), greater scaup (*Aythya marila*), common eider (*Somateria mollissima*), and common and Wilson’s snipe (*Gallinago gallinago* – *G. delicata*). These runner-up models occurred between split-with-migration models and secondary-contact models (Table S3). For demographic analyses we chose the best-fit model to be the one with the lowest AIC value, a ΔAIC = 0, and a weighted AIC = 1 (Tables S3, S4). The best-fit models were: split with symmetric migration (Fig. S1C) for northern pintail (*Anas acuta*), mallard (*Anas platyrhynchos*), and the greater scaup (*Aythya marila*); split with bidirectional (asymmetric) migration (Fig. S1D) in common eider (*Somateria mollissima*), and the common and Wilson’s snipe (*Gallinago gallinago* - *G. delicata*) contrast; secondary contact with symmetric migration (Fig. S1G) in long-tailed ducks (*Clangula hyemalis*); and secondary contact with bidirectional migration (Fig. S1H) for green-winged teal (*Anas crecca crecca – A. c. carolinensis*) and the Eurasian and American wigeon (*Mareca penelope – M. americana*) contrast. From these best-fit models, the raw demographic parameter output (Table S5) was used to calculate a biological estimate of the long-term average rates of gene flow (Table S6).

Estimates for gene flow between Eurasian and North American vector species showed substantial levels of variation in movement (gene flow) between continental populations (Table 1). Values of individuals per generation varied across three orders of magnitude (Table 1). The dabbling ducks—northern pintail (*Anas acuta*), mallard (*Anas platyrhynchos*), Eurasian-American wigeon (*Mareca penelope – M. americana*), and green-winged teal (*Anas crecca crecca – A. c. carolinensis*)—showed the greatest magnitude of movements. Of these, northern pintail (*Anas acuta*) had the largest amount of movement (upwards of ∼100-300 individuals/generation). The diving and sea ducks—long-tailed duck (*Clangula hyemalis*), greater scaup (*Aythya marila*), and common eider (*Somateria mollissima*)—showed a wide range in their levels of movement. Of these ducks, the greatest magnitude of intercontinental movement occurred in long-tailed ducks (*Clangula hyemalis*; ∼24-87 individuals/generation), while the lowest levels occurred in the common eider (∼1-2 individuals/generation; Table 1). Our comparison of common and Wilson’s snipe (*Gallinago gallinago – G. delicata*) showed low levels of movement, with less than 1 individual/generation (Table 1).

**Table 1.**
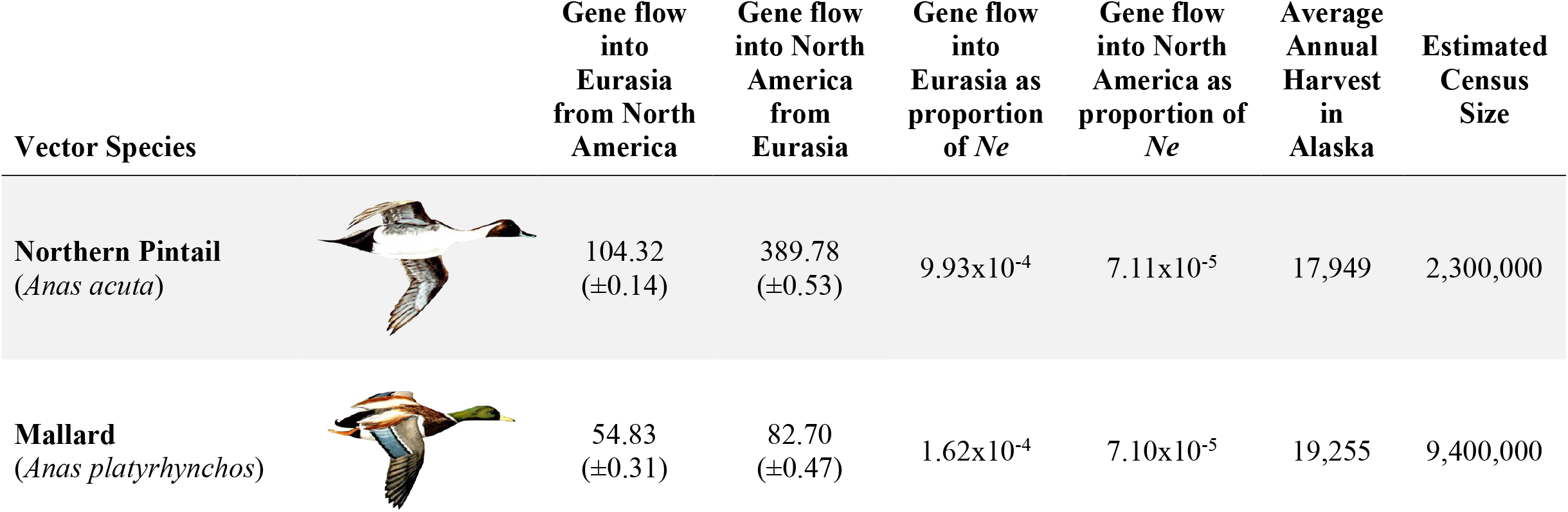
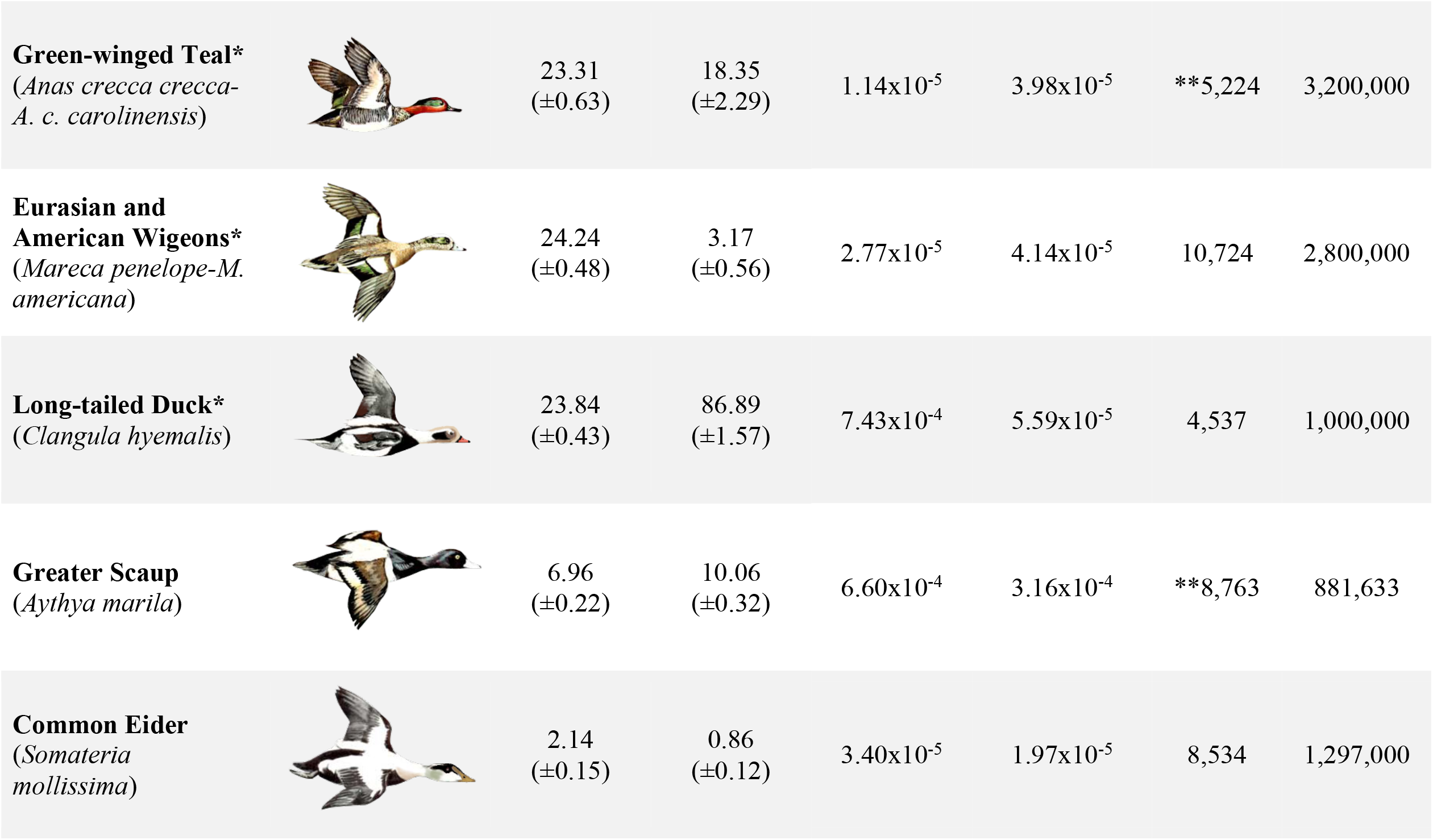
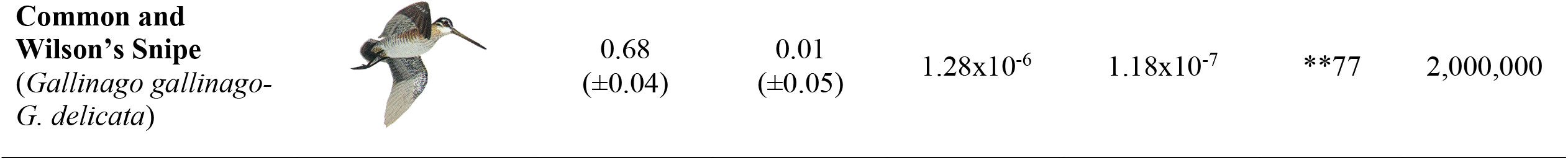
Avian influenza (AI) vector species of birds with estimated rates of gene flow between Eurasian and North American populations (individuals/generation), movement between Eurasian and North American populations as a proportion of effective population size that is experiencing the influx (continent-based *N*_*e*_), average number of individuals harvested in Alaska (across years: 2016, 2017, and 2018), and estimated census size in across North America. Illustrations were obtained from Hines (1963) and National Geographic Society (1999). *Gene flow results from McLaughlin et al. (2020) **Individuals categorized as “Teal” were used for green-winged teal comparison, “Scaup” for greater scaup, and “Small Shorebird” for common and Wilson’s snipe.

Estimates of gene flow between Eurasian and North American populations as a proportion of effective population size (*N*_*e*_) showed the proportion to be small across all species (Table 1). Of all these vector species, greater scaup (*Aythya marila*) had the highest proportion of movement into North America from Eurasia. The northern pintail (*Anas acuta*) had the highest proportion of movement into Eurasia from North America.

### 1.4.2 Relative risks of AI host species

Gene flow as a proportion of *N*_*e*_ is conservative for contemporary populations, and scaling these values to census population sizes and annual harvest rates should result in more useful numbers for risk assessment. We did these calculations using the data from Table 1 to rank vector species in decreasing order of relative risk of intercontinental AI movement, both within the natural system (gene flow scaled to census size) and in the context of subsistence use (gene flow scaled to harvest rates; Table 2). The top three species in each context (though in slightly different order between each context) were greater scaup (*Aythya marila*), mallard (*Anas platyrhynchos*), and northern pintail (*Anas acuta*; Table 2). Other species, e.g., green-winged teal (*Anas crecca crecca - carolinensis*), Eurasian and American wigeons (*Mareca penelope - americana*), and long-tailed duck (*Clangula hyemalis*), were intermixed in their rankings. Finally, common eiders (*Somateria mollissima*) and common and Wilson’s snipe (*Gallinago gallinago – G. delicata*) were equivalently ranked in the two contexts (natural system and subsistence use) and had the lowest relative risk (Table 2).

**Table 2.**
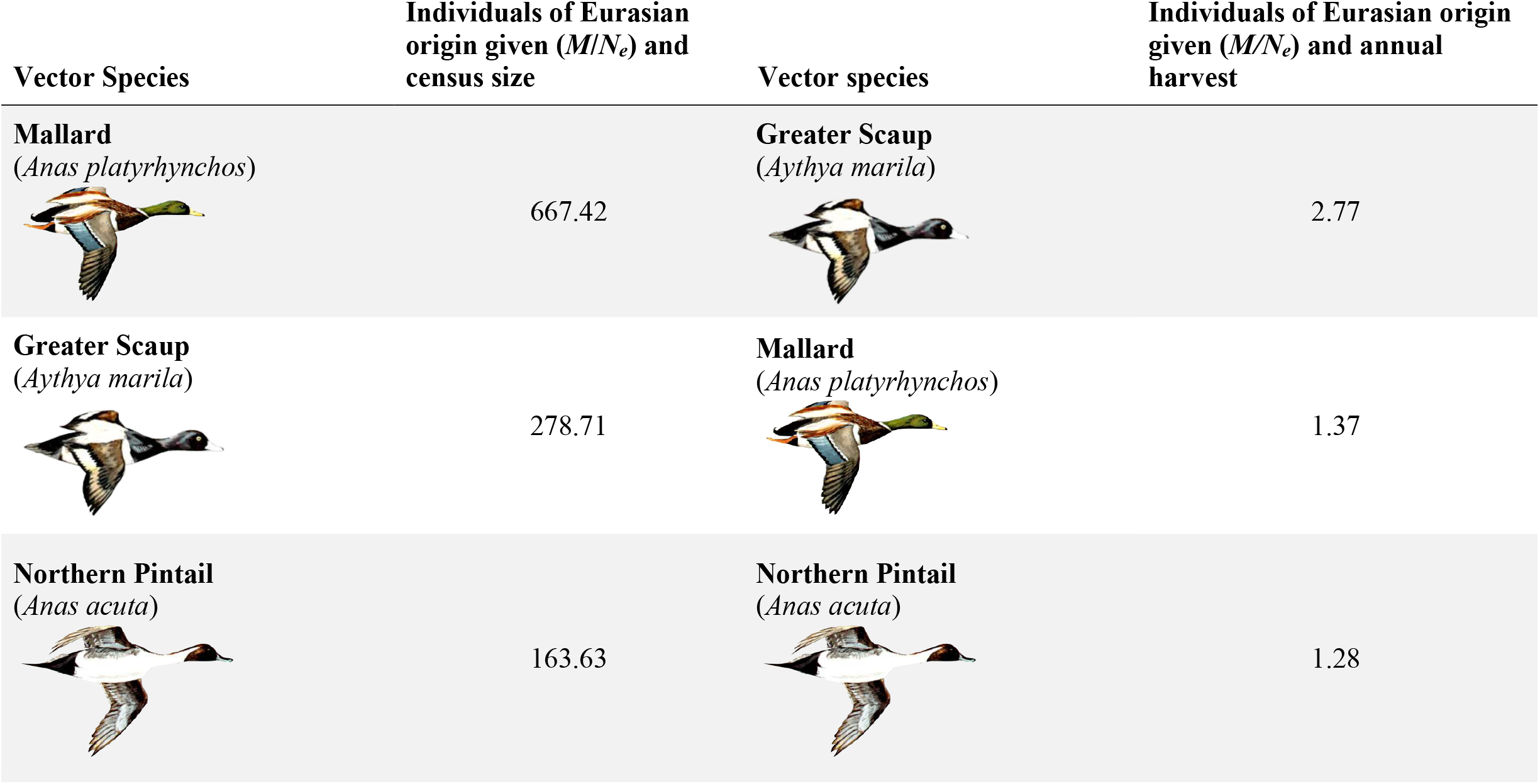
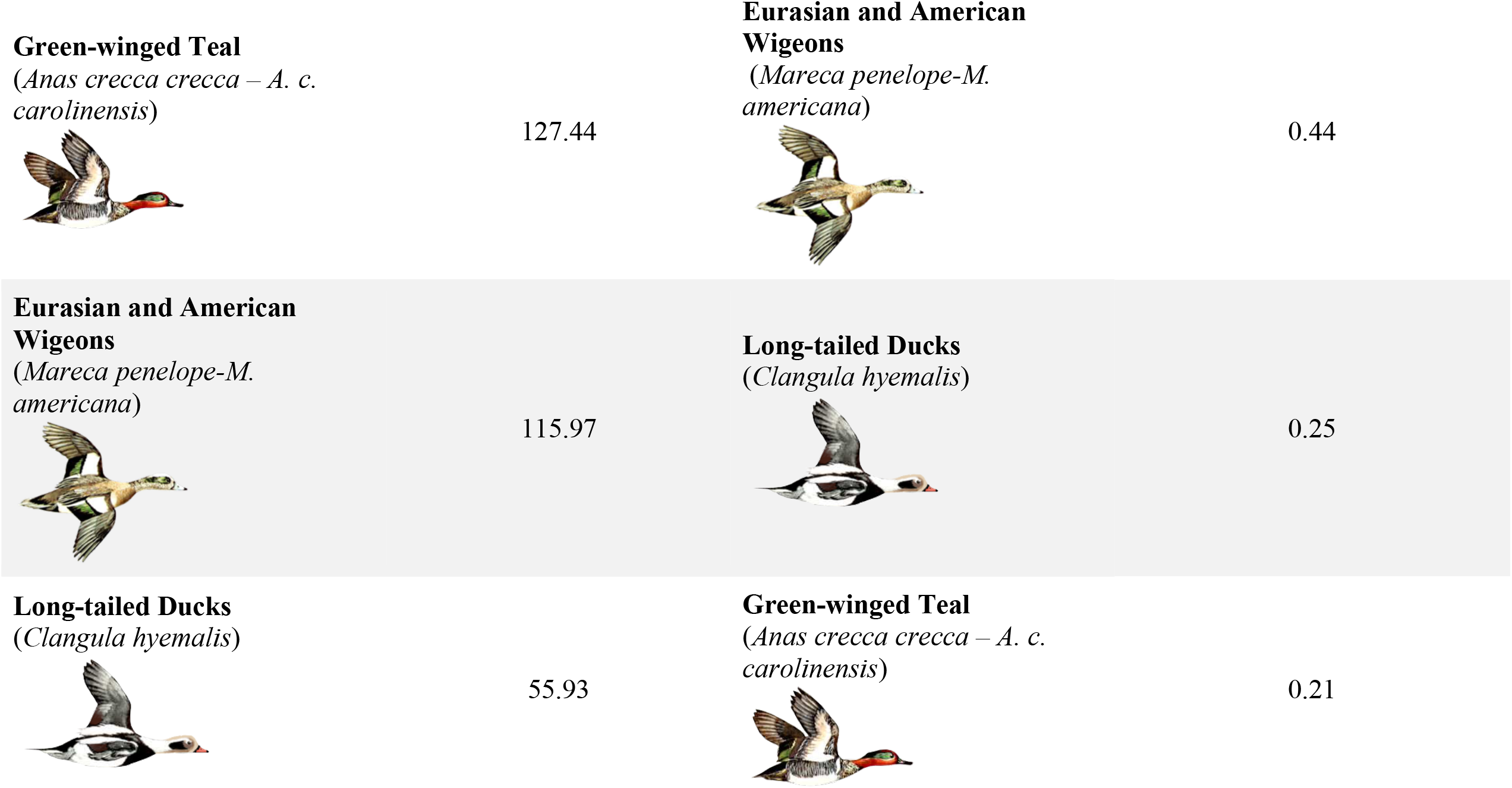
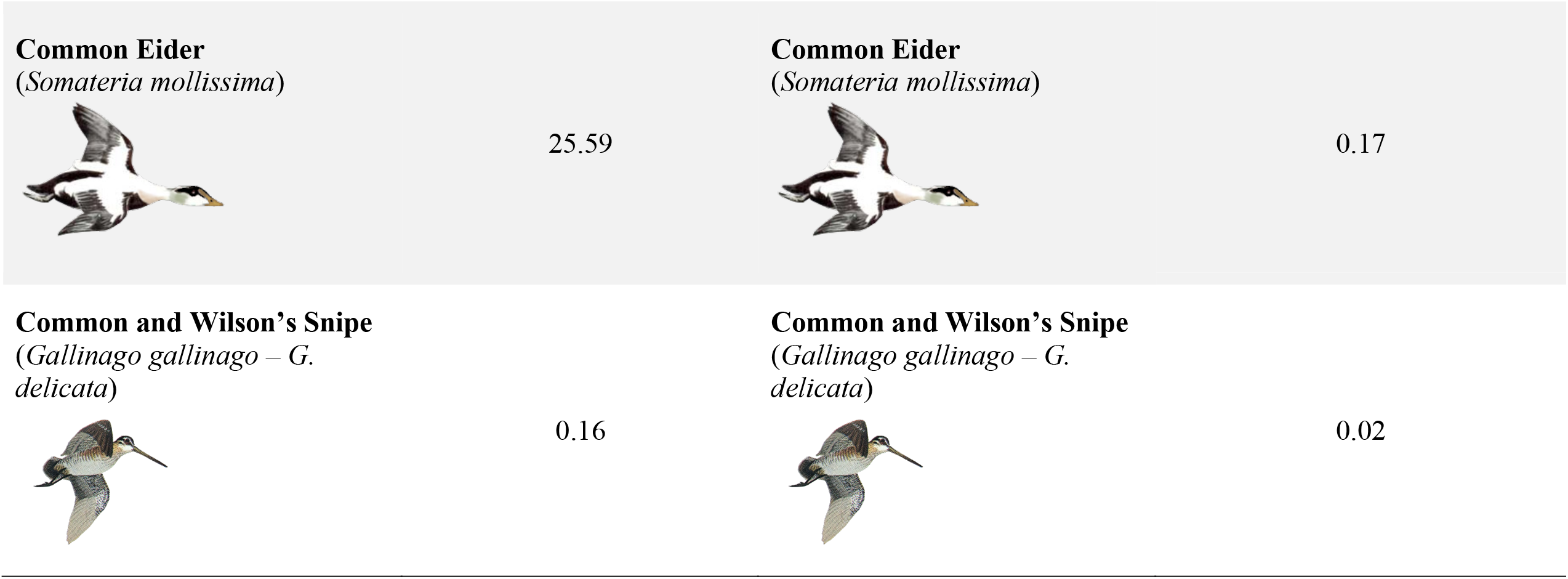
Avian influenza (AI) vector species of birds ranked in decreasing order of their relative risk of AI transmission to North American populations (left column) and to subsistence users (right column). Each entry applies the gene flow estimate from Table 1 to estimate the proportion of census size (left column) or average annual subsistence harvest in Alaska (right column). Illustrations from Hines (1963) and National Geographic Society (1999).

The average annual harvest rates in Alaska (Table 1) show that three taxa, mallard (*Anas platyrhynchos*), northern pintail (*Anas acuta*), and Eurasian and American wigeons (*Mareca penelope - americana*), have the highest rates of harvest. When considering proportions of Eurasian-origin birds, two of these species, the mallard and northern pintail, are ranked numbers two and three (respectively) in relative risk for Asian-origin AI exposure (Table 2). Greater scaup (*Aythya marila*) has only a moderate level of harvest (Table 1), but when considering the estimated proportion of Asian-origin individuals, it had the highest risk of causing potential Asian-origin AI exposure to subsistence users (Table 2).

## 1.5 Discussion

Our study found that levels of intercontinental gene flow in waterbird AI vector species is highly variable. These results provide a solid quantitative framework for among-species contrasts in this AI risk assessment context. We show how population genomics methods can be applied to fill knowledge gaps to help understand the ecology of intercontinental pathogen movements, and to help subsistence users mitigate potential Eurasian-origin AI exposure. Among our sampled species, dabbling ducks overall had the highest rates of intercontinental movement as inferred from long-term average levels of gene flow. Variation in gene flow rates spanning orders of magnitude among AI vector species makes it clear that all do not pose similar risks of intercontinental virus movement (Table 1), and we are able to provide relative risk rankings among the species studied (Table 2). We note that the variation in movement among vector species could be due to variation in factors such as different life-history characteristics, patterns of dispersal, and behavioral tendencies. Assessments of population connectivity are particularly difficult for species that nest at high latitudes, as they often have large distributions across the annual cycle that span remote (uninhabited) landscapes, further reducing the detectability of dispersal events and the ability to evaluate the magnitudes of movements occurring (Sonsthagen et al., 2011; Talbot et al., 2015; Sonsthagen et al., 2019). Overall, our results showed that these vector species appear to vary by three orders of magnitude in the bird-to-bird natural system and by two orders of magnitude in the human-to-bird subsistence system (Tables 1, 2).

Contrasting rates of gene flow, census population sizes, and harvest should enable a more fine-tuned approach to AI risk mitigation, monitoring, and surveillance. However, this approach does have limitations. First, we focused on the host movement system rather than on the viruses themselves. Infection rates are highly variable and are dependent on species, year, seasonality, and geographic location (Webster et al., 1992; Dushoff et al., 2004; Altizer et al., 2006; Winker et al., 2007). Because infection rates are highly variable and are dependent on many factors, we did not incorporate them into our assessment of risk ranking. We feel that until those variables can be better developed and understood in this region, the basic attributes of the natural delivery system (e.g., gene flow rates, census sizes) provide a useful initial baseline for relative risk.

Gene flow is a measure (in evolutionary time) of contact between populations, and it is not equivalent to annual intercontinental movements of individuals. It does, however, provide an effective long-term metric, comparable among lineages, on which to base intercontinental AI movement risk assessments. Our estimates of gene flow are based on long-term *N*_*e*_, and scaling up to census sizes is not straightforward. Long-term effective population sizes are generally smaller than modern census sizes, but among species the two are only loosely correlated (Roman, 2003). So while we have confidence that the relative rates of gene flow between populations are directly comparable among lineages across evolutionary time, converting those values to numbers of individuals given current census sizes (which themselves are only estimates, e.g., Barclay, 2012) represents an approximation. In addition, rates of gene flow can be affected by partial reproductive isolation between lineages, which would make them lower relative to actual movements. Furthermore, gene flow is an imperfect proxy for AI virus transmission because the latter can occur without host reproduction and can easily occur across species, depending on varying levels of host resistance to AI viruses. Thus, while our analyses provide a robust contrast among co-distributed lineages over the long term in an evolutionary context, annual variation is present and can only be approximated by our scaling to census sizes and harvest rates to estimate risk, and gene flow is not a perfect proxy for virus transmission. Future work could involve pairing the results of this study with AI infection rates in each species for region, season, and year when such information becomes available.

Our study provides important details about the vector landscape through which AI viruses must navigate to move intercontinentally in this region. By quantifying the movements of vector species in terms of levels of gene flow and long-term effective population size (*N*_*e*_), we produce a robust metric enabling direct comparison among lineages that move intercontinentally in Beringia. By scaling these long-term movement rates to census sizes and harvest rates in Alaska (Table 1), we were able to estimate the relative risk that each species poses for Eurasian-origin AI both in the natural virus delivery system and in the Alaska subsistence harvest (Table 2). These results give subsistence hunters new information that can be used to choose to harvest species with a lower level of intercontinental movement that pose a lower risk of Eurasian-origin AI exposure, rather than harvest species that might pose a higher risk (Fig. 2). Likewise, prioritizing AI surveillance in species with high levels of intercontinental movement might be useful. Our baseline estimates of host-specific movements of AI vector species can be used to model, monitor, and mitigate AI virus movement in the Beringia region.

**Figure 2.**
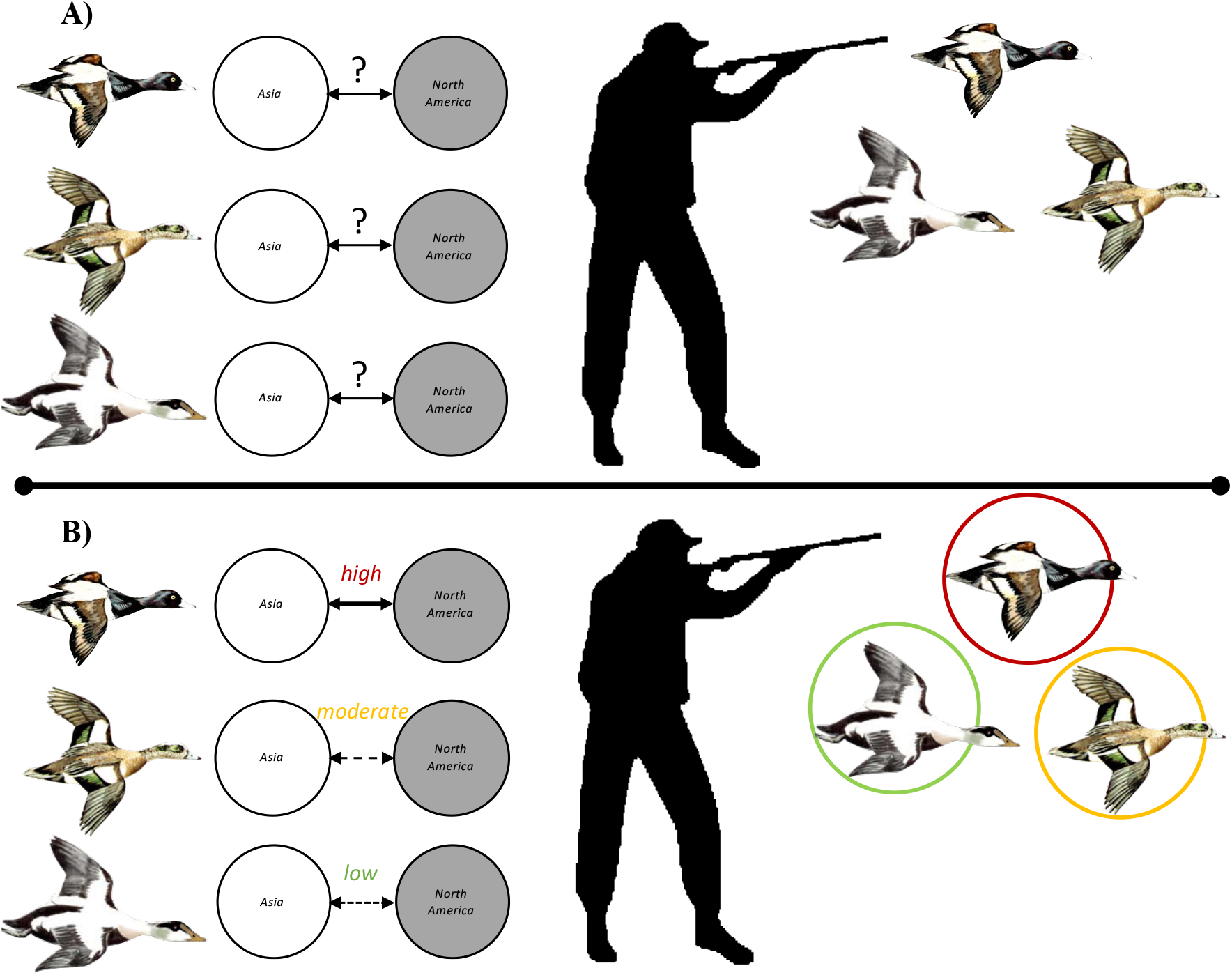
How subsistence hunting could change to mitigate human exposure to Eurasian-origin avian influenza (AI). Species with high levels of intercontinental movement might be less desirable to harvest than species with low to moderate levels of intercontinental movement. **A)** Current understanding: unknown levels of intercontinental movement make it impossible to determine which species pose a high risk of possible Eurasian-origin AI exposure. **B)** This study shows intercontinental movements to be high in greater scaup (red), moderate in wigeons (yellow), and low in common eider (green) given the proportion of individuals of Eurasian origin given (*M/N*_*e*_) and annual harvest (Table 2). With this information, a subsistence hunter can choose to harvest species with lower intercontinental movements at higher rates to lower human risk of potential Eurasian-origin AI exposure. Illustrations from Hines (1963).

## Acknowledgments

We thank the University of Alaska Museum and the Louisiana State University Museum of Natural Science for loans of tissue specimens and the specimen collectors who made it possible to do this research. We also thank the Kessel Fund for Northern Ornithology and the Friends of Ornithology for financial support, and Brant Faircloth for his input early in the project. This project was funded in part by the the National Science Foundation (DEB-1242267-1242241-1242260), by an Institutional Development Award (IDeA) from the National Institute of General Medical Sciences of the National Institutes of Health under grant number 2P20GM103395, and by the National Institute of General Medical Sciences of the National Institutes of Health under award number RL5GM118990. Naoki Takebayashi and Devin Drown provided helpful comments and feedback on earlier drafts of the manuscript.

## Data Availability Statement

Original sequence data have been deposited in the NCBI Sequence Read Archive (SRA; Table 1.S1; projects PRJNA741698, PRJNA741809, and PRJNA393740).

## Author Contributions

The study was conceived by Kevin Winker, Travis Glenn, and Brant Faircloth. The data were generated by Travis Glenn and Brant Faircloth. The data were analyzed by Fern Spaulding. The first draft was written by Fern Spaulding. All authors contributed to the final version.

## 1.9 Appendix

### Supplementary Information

**Table S1.**
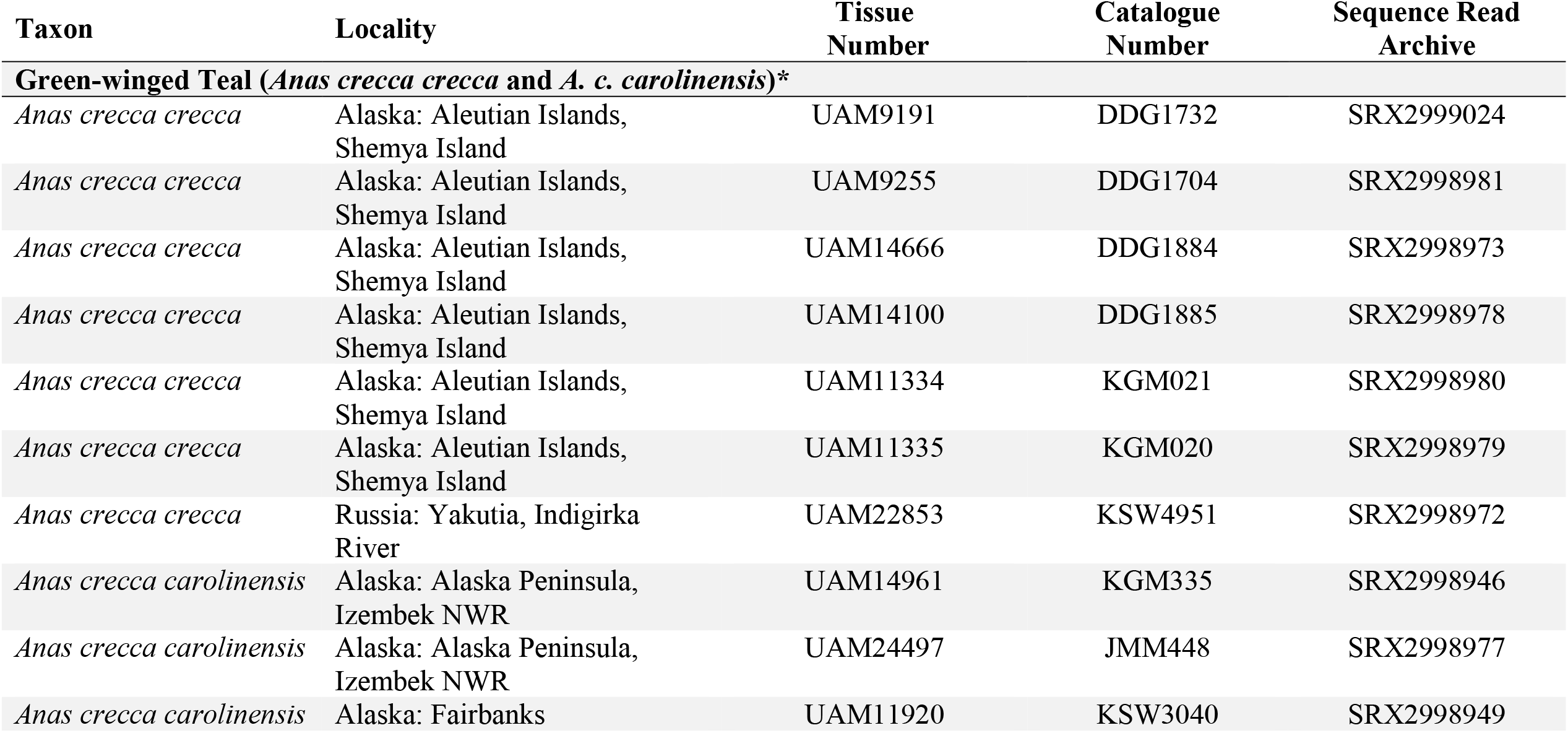
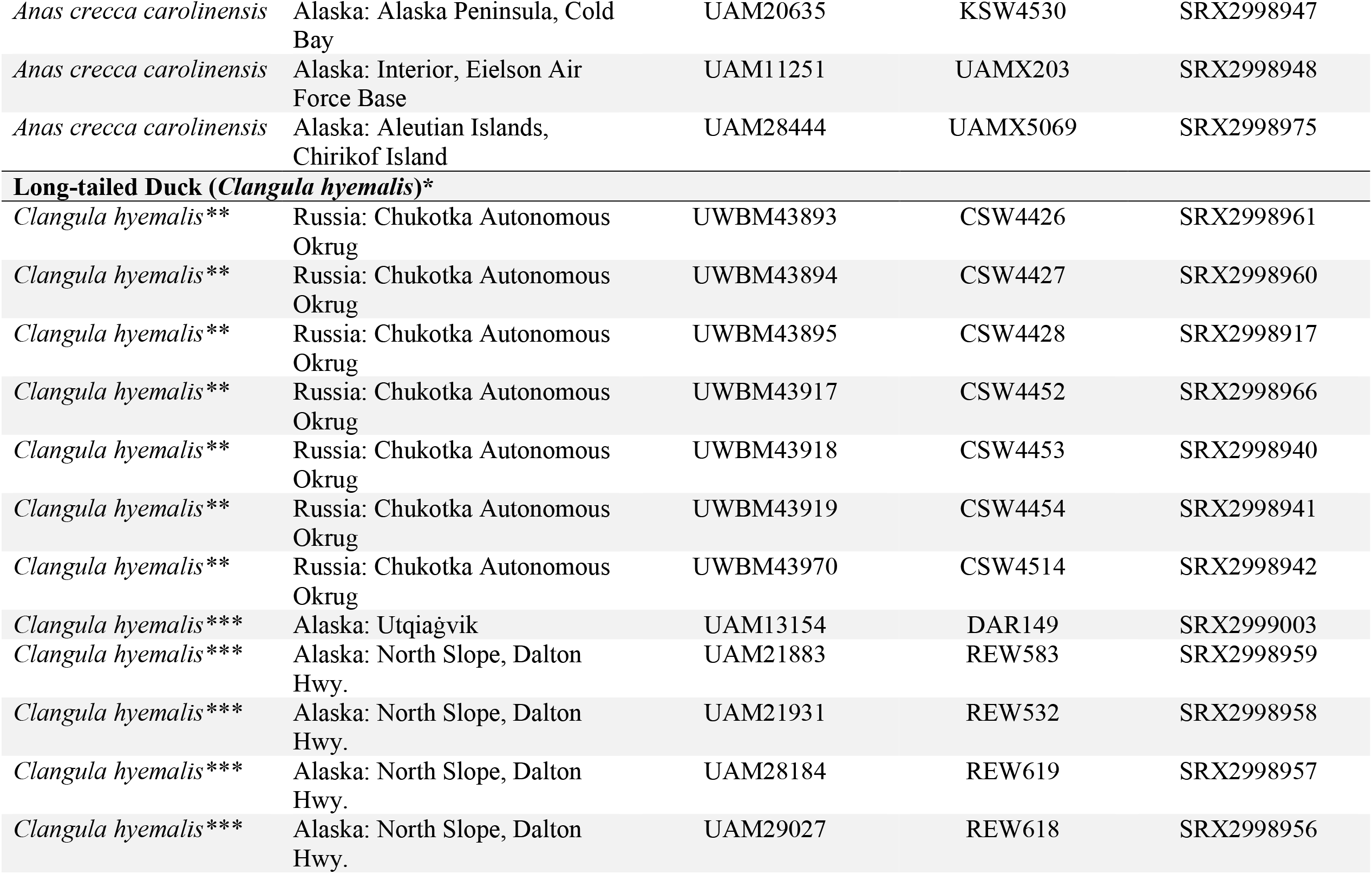
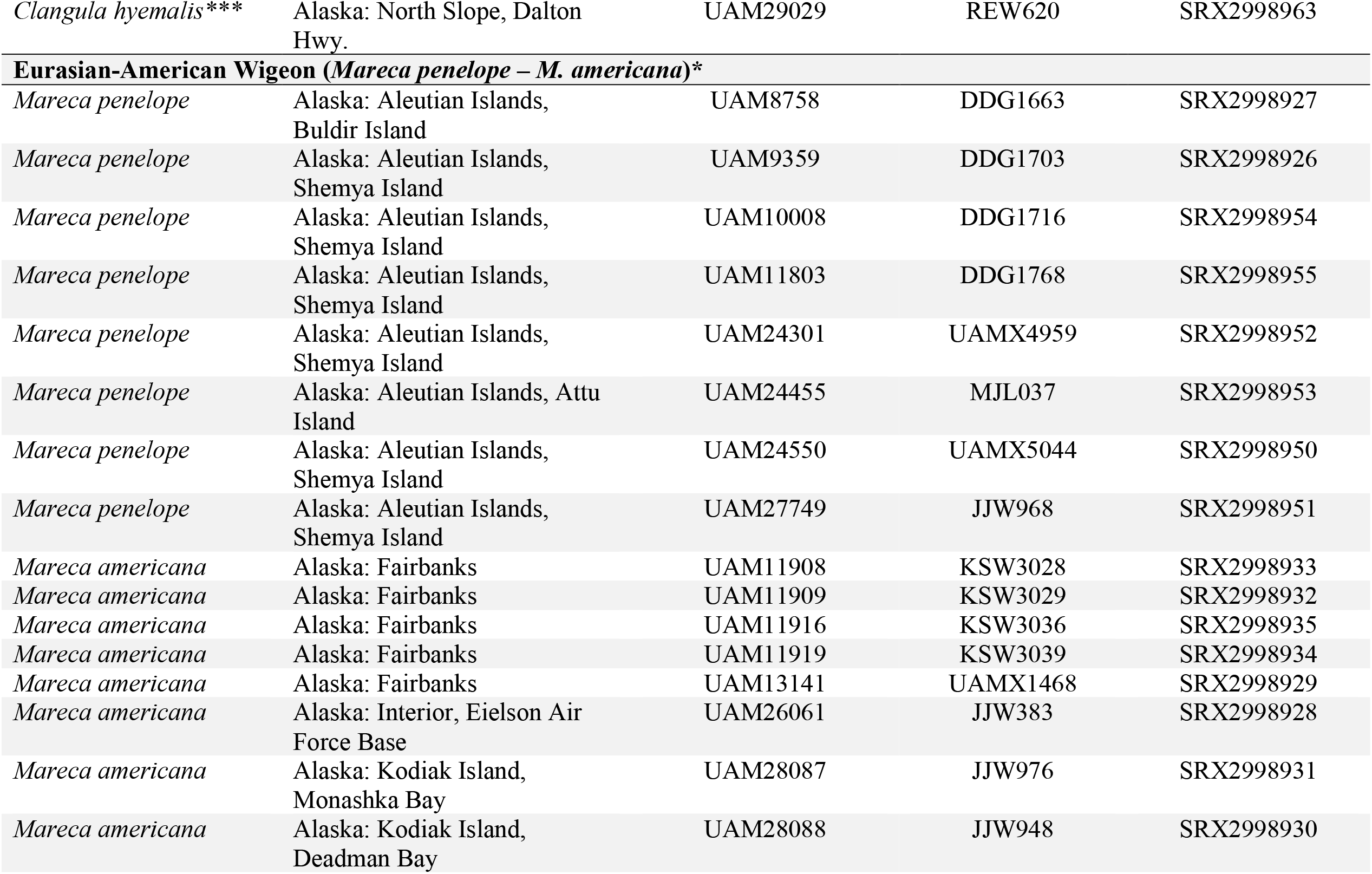
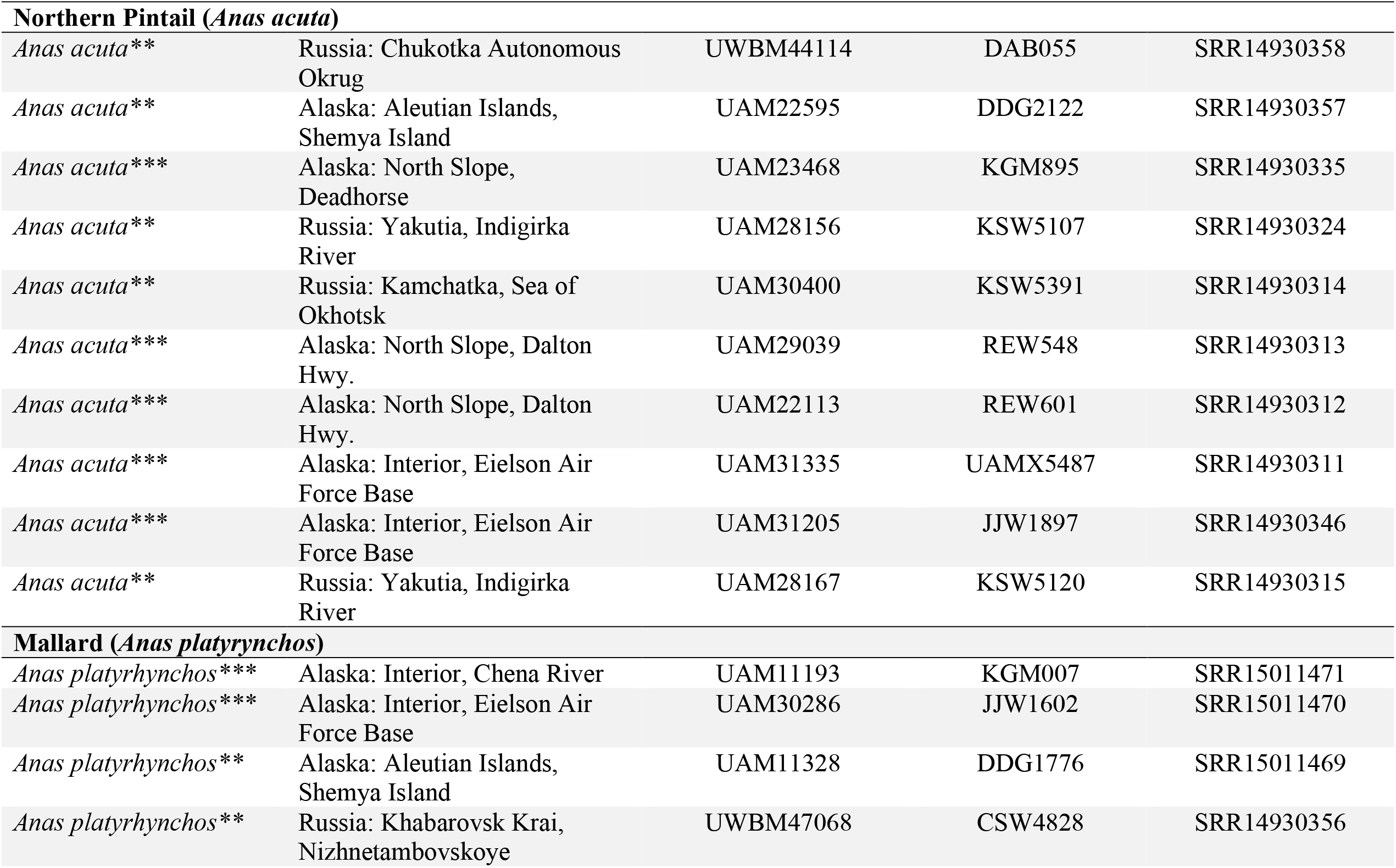
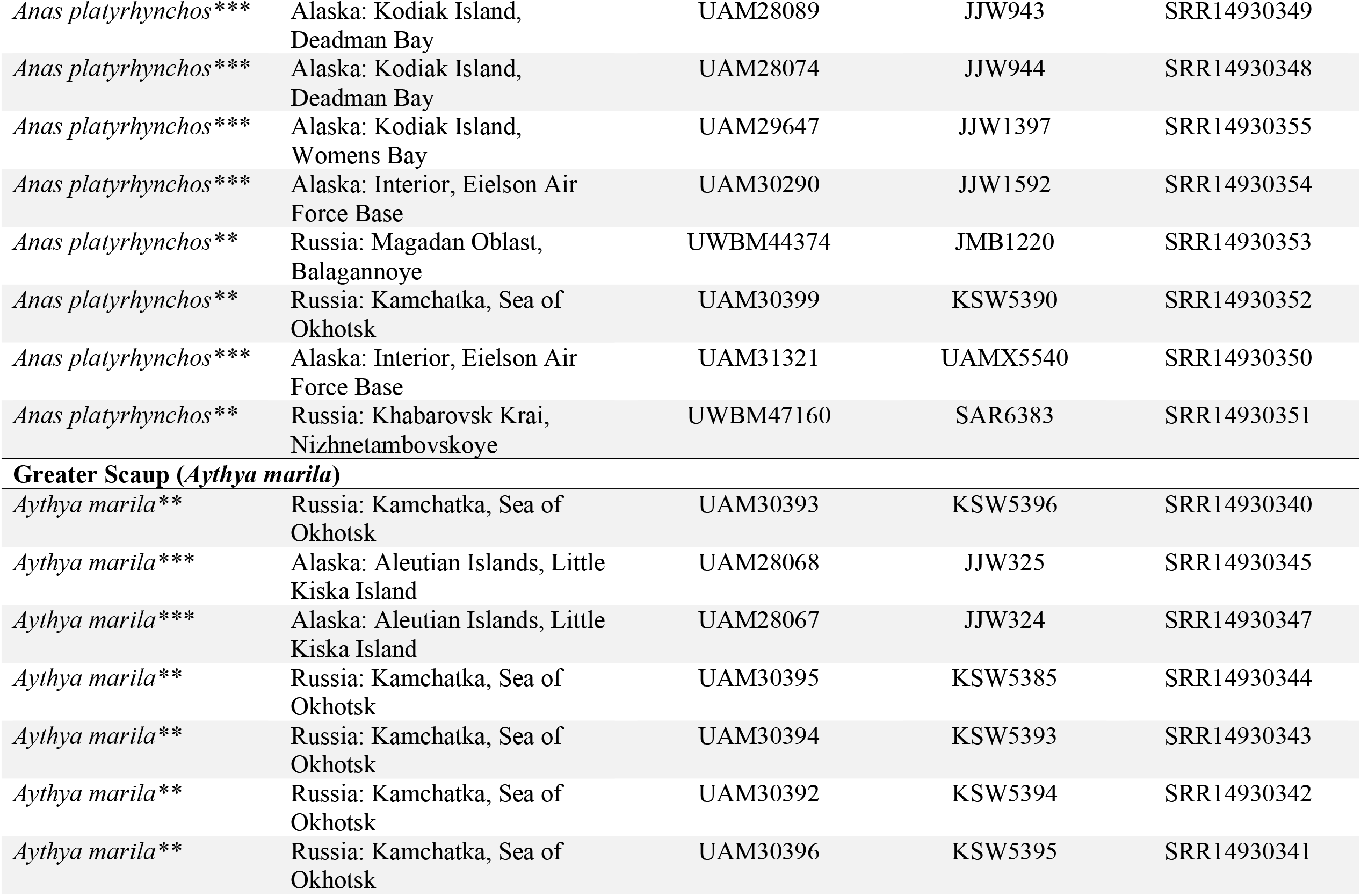
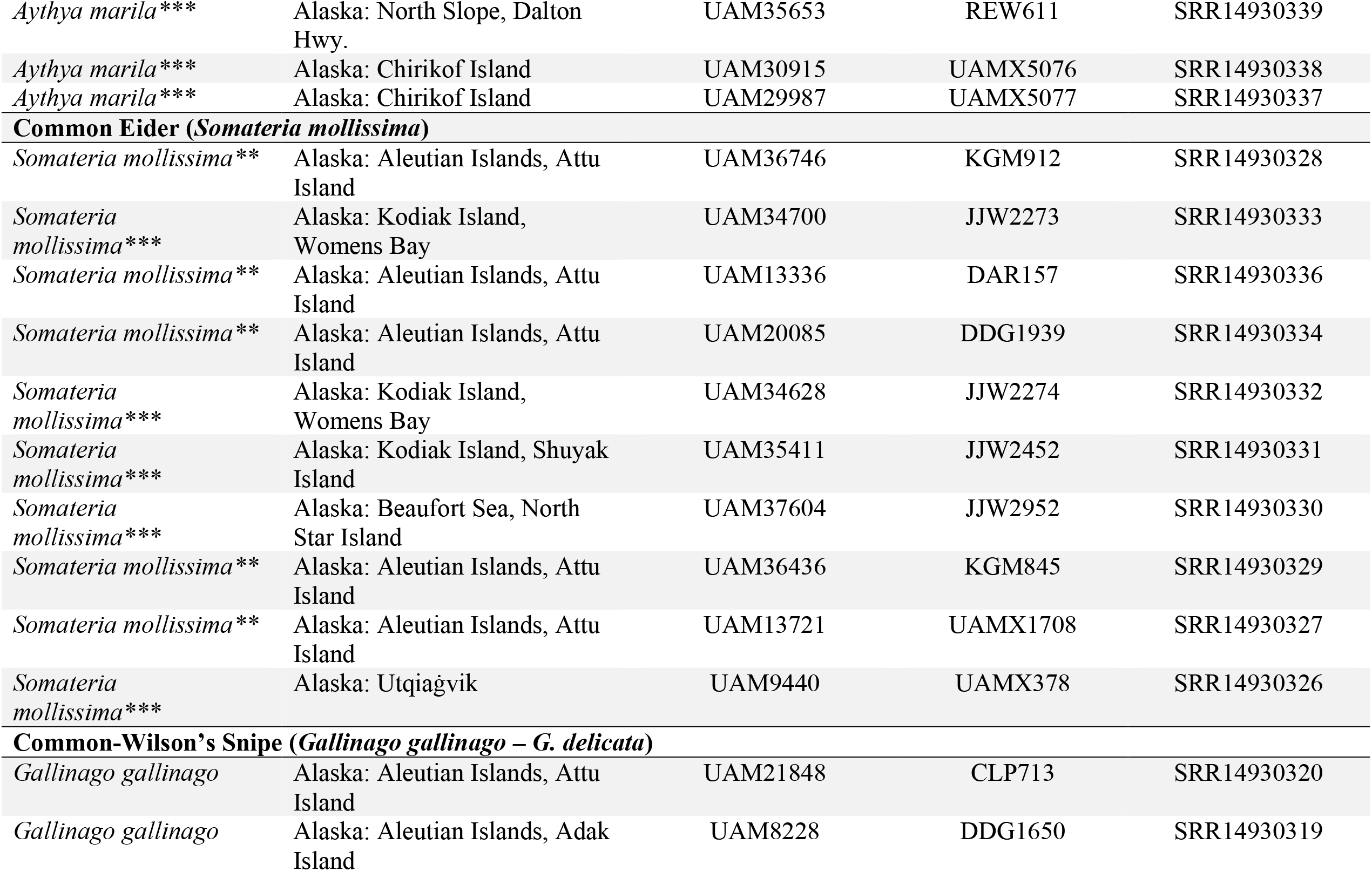
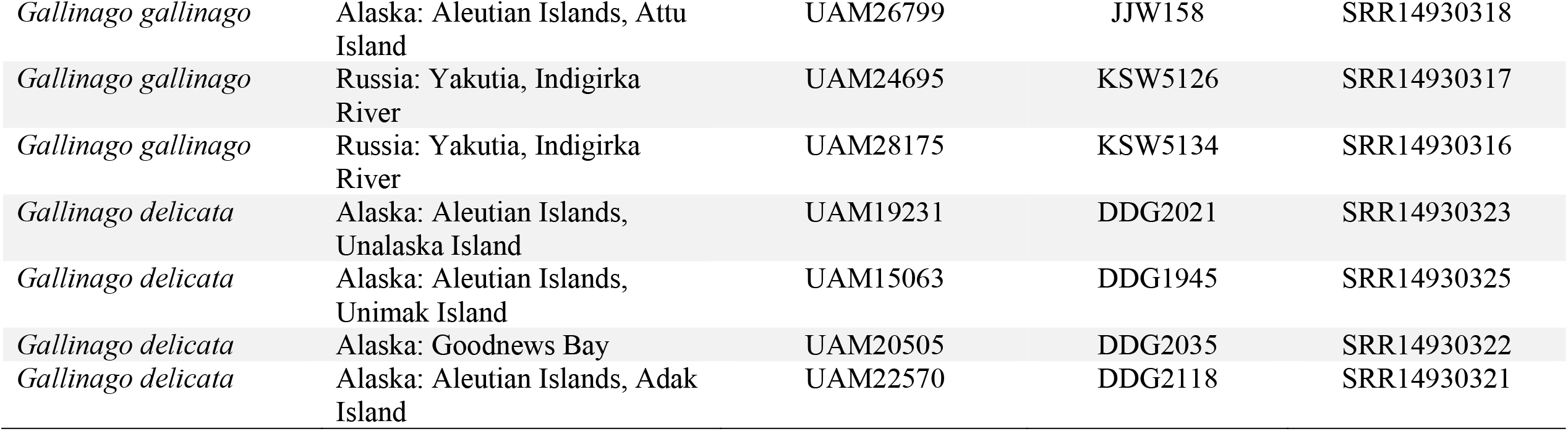
Specimen information of individuals used in this study. Note that UAM = University of Alaska Museum; UWBM = University of Washington Burke Museum. All SRA reads available under project PRJNA741809 and project PRJNA741698. *Individuals from McLaughlin et al. (2020). All SRA reads available under project PRJNA393740. **Individuals considered to be of Eurasian origin (if not classified at the subspecies or species level). ***Individuals considered to be of North American origin (if not classified at the subspecies or species level).

| Taxon | Locality | Tissue Number | Catalogue Number | Sequence Read Archive |
| --- | --- | --- | --- | --- |
| **Green-winged Teal (*Anas crecca crecca* and *A. c. carolinensis*)\*** |  |  |  |  |
| *Anas crecca crecca* | Alaska: Aleutian Islands, Shemya Island | UAM9191 | DDG1732 | SRX2999024 |
| *Anas crecca crecca* | Alaska: Aleutian Islands, Shemya Island | UAM9255 | DDG1704 | SRX2998981 |
| *Anas crecca crecca* | Alaska: Aleutian Islands, Shemya Island | UAM14666 | DDG1884 | SRX2998973 |
| *Anas crecca crecca* | Alaska: Aleutian Islands, Shemya Island | UAM14100 | DDG1885 | SRX2998978 |
| *Anas crecca crecca* | Alaska: Aleutian Islands, Shemya Island | UAM11334 | KGM021 | SRX2998980 |
| *Anas crecca crecca* | Alaska: Aleutian Islands, Shemya Island | UAM11335 | KGM020 | SRX2998979 |
| *Anas crecca crecca* | Russia: Yakutia, Indigirka River | UAM22853 | KSW4951 | SRX2998972 |
| *Anas crecca carolinensis* | Alaska: Alaska Peninsula, Izembek NWR | UAM14961 | KGM335 | SRX2998946 |
| *Anas crecca carolinensis* | Alaska: Alaska Peninsula, Izembek NWR | UAM24497 | JMM448 | SRX2998977 |
| *Anas crecca carolinensis* | Alaska: Fairbanks | UAM11920 | KSW3040 | SRX2998949 |
| *Anas crecca carolinensis* | Alaska: Alaska Peninsula, Cold Bay | UAM20635 | KSW4530 | SRX2998947 |
| *Anas crecca carolinensis* | Alaska: Interior, Eielson Air Force Base | UAM11251 | UAMX203 | SRX2998948 |
| *Anas crecca carolinensis* | Alaska: Aleutian Islands, Chirikof Island | UAM28444 | UAMX5069 | SRX2998975 |
| **Long-tailed Duck (*Clangula hyemalis*)\*** |  |  |  |  |
| *Clangula hyemalis* ** | Russia: Chukotka Autonomous Okrug | UWBM43893 | CSW4426 | SRX2998961 |
| *Clangula hyemalis* ** | Russia: Chukotka Autonomous Okrug | UWBM43894 | CSW4427 | SRX2998960 |
| *Clangula hyemalis* ** | Russia: Chukotka Autonomous Okrug | UWBM43895 | CSW4428 | SRX2998917 |
| *Clangula hyemalis* ** | Russia: Chukotka Autonomous Okrug | UWBM43917 | CSW4452 | SRX2998966 |
| *Clangula hyemalis* ** | Russia: Chukotka Autonomous Okrug | UWBM43918 | CSW4453 | SRX2998940 |
| *Clangula hyemalis* ** | Russia: Chukotka Autonomous Okrug | UWBM43919 | CSW4454 | SRX2998941 |
| *Clangula hyemalis* ** | Russia: Chukotka Autonomous Okrug | UWBM43970 | CSW4514 | SRX2998942 |
| *Clangula hyemalis* *** | Alaska: Utqiagvik | UAM13154 | DAR149 | SRX2999003 |
| *Clangula hyemalis* *** | Alaska: North Slope, Dalton Hwy. | UAM21883 | REW583 | SRX2998959 |
| *Clangula hyemalis* *** | Alaska: North Slope, Dalton Hwy. | UAM21931 | REW532 | SRX2998958 |
| *Clangula hyemalis* *** | Alaska: North Slope, Dalton Hwy. | UAM28184 | REW619 | SRX2998957 |
| *Clangula hyemalis* *** | Alaska: North Slope, Dalton Hwy. | UAM29027 | REW618 | SRX2998956 |
| *Clangula hyemalis* *** | Alaska: North Slope, Dalton Hwy. | UAM29029 | REW620 | SRX2998963 |
| **Eurasian-American Wigeon (*Mareca penelope* – *M. americana*)\*** |  |  |  |  |
| *Mareca penelope* | Alaska: Aleutian Islands, Buldir Island | UAM8758 | DDG1663 | SRX2998927 |
| *Mareca penelope* | Alaska: Aleutian Islands, Shemya Island | UAM9359 | DDG1703 | SRX2998926 |
| *Mareca penelope* | Alaska: Aleutian Islands, Shemya Island | UAM10008 | DDG1716 | SRX2998954 |
| *Mareca penelope* | Alaska: Aleutian Islands, Shemya Island | UAM11803 | DDG1768 | SRX2998955 |
| *Mareca penelope* | Alaska: Aleutian Islands, Shemya Island | UAM24301 | UAMX4959 | SRX2998952 |
| *Mareca penelope* | Alaska: Aleutian Islands, Attu Island | UAM24455 | MJL037 | SRX2998953 |
| *Mareca penelope* | Alaska: Aleutian Islands, Shemya Island | UAM24550 | UAMX5044 | SRX2998950 |
| *Mareca penelope* | Alaska: Aleutian Islands, Shemya Island | UAM27749 | JJW968 | SRX2998951 |
| *Mareca americana* | Alaska: Fairbanks | UAM11908 | KSW3028 | SRX2998933 |
| *Mareca americana* | Alaska: Fairbanks | UAM11909 | KSW3029 | SRX2998932 |
| *Mareca americana* | Alaska: Fairbanks | UAM11916 | KSW3036 | SRX2998935 |
| *Mareca americana* | Alaska: Fairbanks | UAM11919 | KSW3039 | SRX2998934 |
| *Mareca americana* | Alaska: Fairbanks | UAM13141 | UAMX1468 | SRX2998929 |
| *Mareca americana* | Alaska: Interior, Eielson Air Force Base | UAM26061 | JJW383 | SRX2998928 |
| *Mareca americana* | Alaska: Kodiak Island, Monashka Bay | UAM28087 | JJW976 | SRX2998931 |
| *Mareca americana* | Alaska: Kodiak Island, Deadman Bay | UAM28088 | JJW948 | SRX2998930 |

| **Northern Pintail (*Anas acuta*)** |  |  |  |  |
| --- | --- | --- | --- | --- |
| *Anas acuta* ** | Russia: Chukotka Autonomous Okrug | UWBM44114 | DAB055 | SRR14930358 |
| *Anas acuta* ** | Alaska: Aleutian Islands, Shemya Island | UAM22595 | DDG2122 | SRR14930357 |
| *Anas acuta* *** | Alaska: North Slope, Deadhorse | UAM23468 | KGM895 | SRR14930335 |
| *Anas acuta* ** | Russia: Yakutia, Indigirka River | UAM28156 | KSW5107 | SRR14930324 |
| *Anas acuta* ** | Russia: Kamchatka, Sea of Okhotsk | UAM30400 | KSW5391 | SRR14930314 |
| *Anas acuta* *** | Alaska: North Slope, Dalton Hwy. | UAM29039 | REW548 | SRR14930313 |
| *Anas acuta* *** | Alaska: North Slope, Dalton Hwy. | UAM22113 | REW601 | SRR14930312 |
| *Anas acuta* *** | Alaska: Interior, Eielson Air Force Base | UAM31335 | UAMX5487 | SRR14930311 |
| *Anas acuta* *** | Alaska: Interior, Eielson Air Force Base | UAM31205 | JJW1897 | SRR14930346 |
| *Anas acuta* ** | Russia: Yakutia, Indigirka River | UAM28167 | KSW5120 | SRR14930315 |
| **Mallard (*Anas platyrhynchos*)** |  |  |  |  |
| *Anas platyrhynchos* *** | Alaska: Interior, Chena River | UAM11193 | KGM007 | SRR15011471 |
| *Anas platyrhynchos* *** | Alaska: Interior, Eielson Air Force Base | UAM30286 | JJW1602 | SRR15011470 |
| *Anas platyrhynchos* ** | Alaska: Aleutian Islands, Shemya Island | UAM11328 | DDG1776 | SRR15011469 |
| *Anas platyrhynchos* ** | Russia: Khabarovsk Krai, Nizhnetambovskoye | UWBM47068 | CSW4828 | SRR14930356 |
| *Anas platyrhynchos* *** | Alaska: Kodiak Island,<br>Deadman Bay | UAM28089 | JJW943 | SRR14930349 |
| *Anas platyrhynchos* *** | Alaska: Kodiak Island,<br>Deadman Bay | UAM28074 | JJW944 | SRR14930348 |
| *Anas platyrhynchos* *** | Alaska: Kodiak Island,<br>Womens Bay | UAM29647 | JJW1397 | SRR14930355 |
| *Anas platyrhynchos* *** | Alaska: Interior, Eielson Air<br>Force Base | UAM30290 | JJW1592 | SRR14930354 |
| *Anas platyrhynchos* ** | Russia: Magadan Oblast,<br>Balagannoye | UWBM44374 | JMB1220 | SRR14930353 |
| *Anas platyrhynchos* ** | Russia: Kamchatka, Sea of<br>Okhotsk | UAM30399 | KSW5390 | SRR14930352 |
| *Anas platyrhynchos* *** | Alaska: Interior, Eielson Air<br>Force Base | UAM31321 | UAMX5540 | SRR14930350 |
| *Anas platyrhynchos* ** | Russia: Khabarovsk Krai,<br>Nizhnetambovskoye | UWBM47160 | SAR6383 | SRR14930351 |
| **Greater Scaup (*Aythya marila*)** |  |  |  |  |
| *Aythya marila* ** | Russia: Kamchatka, Sea of<br>Okhotsk | UAM30393 | KSW5396 | SRR14930340 |
| *Aythya marila* *** | Alaska: Aleutian Islands, Little<br>Kiska Island | UAM28068 | JJW325 | SRR14930345 |
| *Aythya marila* *** | Alaska: Aleutian Islands, Little<br>Kiska Island | UAM28067 | JJW324 | SRR14930347 |
| *Aythya marila* ** | Russia: Kamchatka, Sea of<br>Okhotsk | UAM30395 | KSW5385 | SRR14930344 |
| *Aythya marila* ** | Russia: Kamchatka, Sea of<br>Okhotsk | UAM30394 | KSW5393 | SRR14930343 |
| *Aythya marila* ** | Russia: Kamchatka, Sea of<br>Okhotsk | UAM30392 | KSW5394 | SRR14930342 |
| *Aythya marila* ** | Russia: Kamchatka, Sea of<br>Okhotsk | UAM30396 | KSW5395 | SRR14930341 |
| *Aythya marila* *** | Alaska: North Slope, Dalton Hwy. | UAM35653 | REW611 | SRR14930339 |
| *Aythya marila* *** | Alaska: Chirikof Island | UAM30915 | UAMX5076 | SRR14930338 |
| *Aythya marila* *** | Alaska: Chirikof Island | UAM29987 | UAMX5077 | SRR14930337 |
| **Common Eider (*Somateria mollissima*)** |  |  |  |  |
| *Somateria mollissima* ** | Alaska: Aleutian Islands, Attu Island | UAM36746 | KGM912 | SRR14930328 |
| *Somateria mollissima* *** | Alaska: Kodiak Island, Womens Bay | UAM34700 | JJW2273 | SRR14930333 |
| *Somateria mollissima* ** | Alaska: Aleutian Islands, Attu Island | UAM13336 | DAR157 | SRR14930336 |
| *Somateria mollissima* ** | Alaska: Aleutian Islands, Attu Island | UAM20085 | DDG1939 | SRR14930334 |
| *Somateria mollissima* *** | Alaska: Kodiak Island, Womens Bay | UAM34628 | JJW2274 | SRR14930332 |
| *Somateria mollissima* *** | Alaska: Kodiak Island, Shuyak Island | UAM35411 | JJW2452 | SRR14930331 |
| *Somateria mollissima* *** | Alaska: Beaufort Sea, North Star Island | UAM37604 | JJW2952 | SRR14930330 |
| *Somateria mollissima* ** | Alaska: Aleutian Islands, Attu Island | UAM36436 | KGM845 | SRR14930329 |
| *Somateria mollissima* ** | Alaska: Aleutian Islands, Attu Island | UAM13721 | UAMX1708 | SRR14930327 |
| *Somateria mollissima* *** | Alaska: Utqiagvik | UAM9440 | UAMX378 | SRR14930326 |
| **Common-Wilson's Snipe (*Gallinago gallinago* – *G. delicata*)** |  |  |  |  |
| *Gallinago gallinago* | Alaska: Aleutian Islands, Attu Island | UAM21848 | CLP713 | SRR14930320 |
| *Gallinago gallinago* | Alaska: Aleutian Islands, Adak Island | UAM8228 | DDG1650 | SRR14930319 |
| *Gallinago gallinago* | Alaska: Aleutian Islands, Attu Island | UAM26799 | JJW158 | SRR14930318 |
| *Gallinago gallinago* | Russia: Yakutia, Indigirka River | UAM24695 | KSW5126 | SRR14930317 |
| *Gallinago gallinago* | Russia: Yakutia, Indigirka River | UAM28175 | KSW5134 | SRR14930316 |
| *Gallinago delicata* | Alaska: Aleutian Islands, Unalaska Island | UAM19231 | DDG2021 | SRR14930323 |
| *Gallinago delicata* | Alaska: Aleutian Islands, Unimak Island | UAM15063 | DDG1945 | SRR14930325 |
| *Gallinago delicata* | Alaska: Goodnews Bay | UAM20505 | DDG2035 | SRR14930322 |
| *Gallinago delicata* | Alaska: Aleutian Islands, Adak Island | UAM22570 | DDG2118 | SRR14930321 |

**Table S2.**
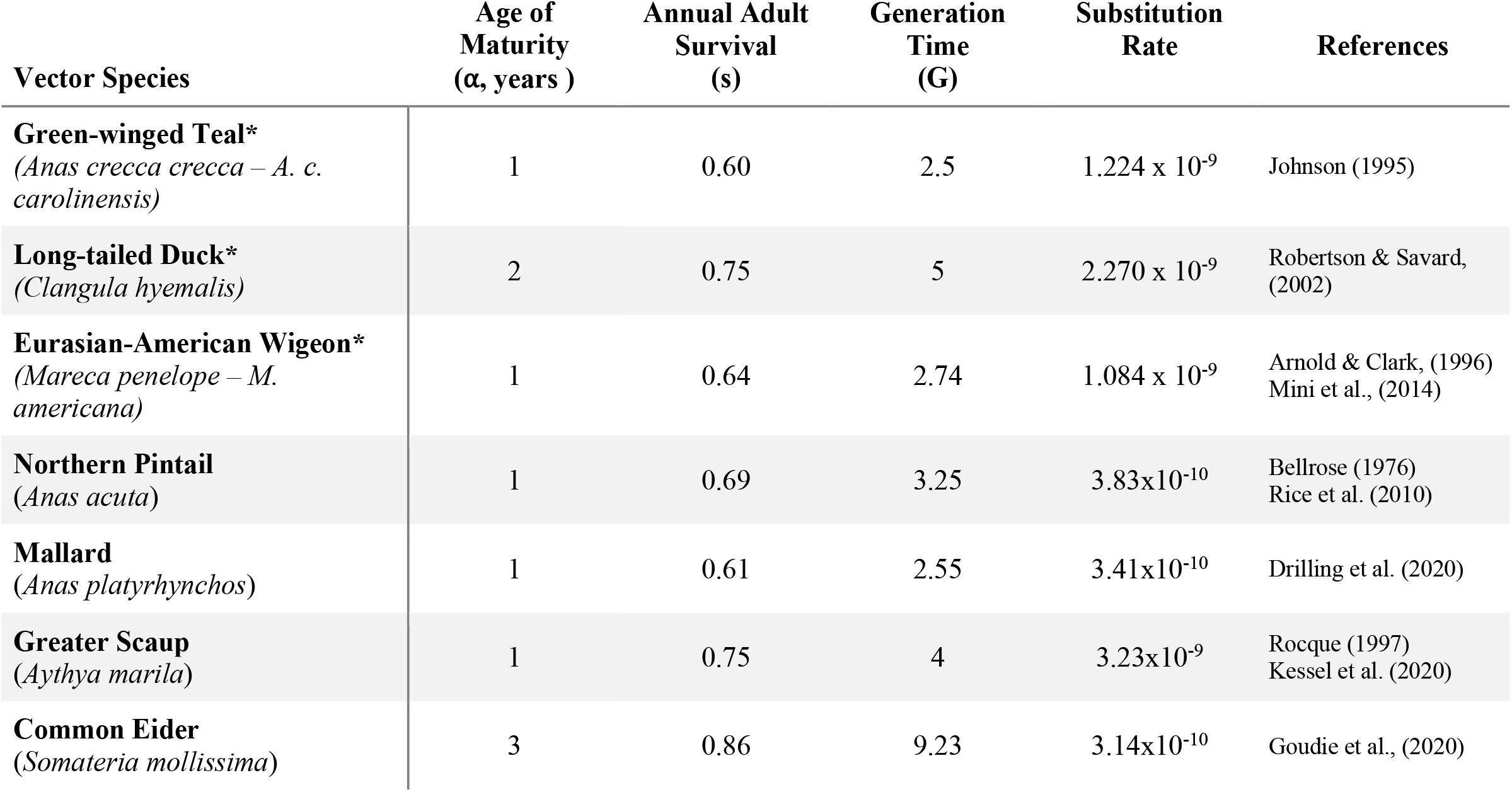
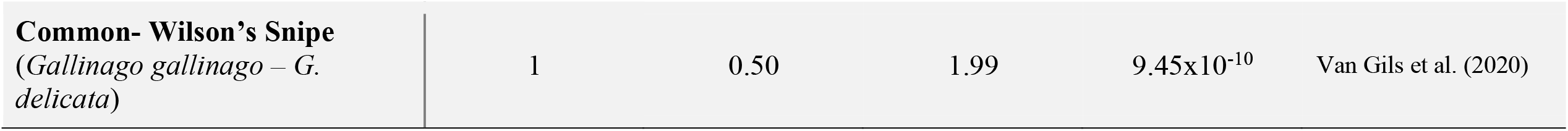
Values used to translate δaδi output into biological estimates of demographic parameters. The equation for generation time *G = α* − *[s/(1-s)]* was obtained from Saether et al. (2005). Substitutions = mutations/site/year. To calculate substitution rates we used fossil-calibrated nodes from Claramunt & Cracraft (2015). In ducks, we used a node within Anseriformes (*Anser cygnoides*; GenBank accession: GCA_000971095.1) with a date of 28.13 Ma. For shorebirds, we used a node within the Charadriiformes (*Charadrius vociferus*; GenBank accession: GCA_000708025.2) with a date of 29.78 Ma. The biological estimate for ancestral population size (*N*_*ref*_), is derived from the output of *Θ* from δaδi, where *Θ* = 4 * *N*_*ref*_ * *substitution rate*. *Results from McLaughlin et al. (2020)

| Vector Species | Age of Maturity ($\alpha$, years ) | Annual Adult Survival (s) | Generation Time (G) | Substitution Rate | References |
| --- | --- | --- | --- | --- | --- |
| **Green-winged Teal\***<br>(*Anas crecca crecca* – *A. c. carolinensis*) | 1 | 0.60 | 2.5 | 1.224 x 10<sup>-9</sup> | Johnson (1995) |
| **Long-tailed Duck\***<br>(*Clangula hyemalis*) | 2 | 0.75 | 5 | 2.270 x 10<sup>-9</sup> | Robertson & Savard, (2002) |
| **Eurasian-American Wigeon\***<br>(*Mareca penelope* – *M. americana*) | 1 | 0.64 | 2.74 | 1.084 x 10<sup>-9</sup> | Arnold & Clark, (1996)<br>Mini et al., (2014) |
| **Northern Pintail**<br>(*Anas acuta*) | 1 | 0.69 | 3.25 | 3.83x10<sup>-10</sup> | Bellrose (1976)<br>Rice et al. (2010) |
| **Mallard**<br>(*Anas platyrhynchos*) | 1 | 0.61 | 2.55 | 3.41x10<sup>-10</sup> | Drilling et al. (2020) |
| **Greater Scaup**<br>(*Aythya marila*) | 1 | 0.75 | 4 | 3.23x10<sup>-9</sup> | Rocque (1997)<br>Kessel et al. (2020) |
| **Common Eider**<br>(*Somateria mollissima*) | 3 | 0.86 | 9.23 | 3.14x10<sup>-10</sup> | Goudie et al., (2020) |
| **Common- Wilson's Snipe**<br>(*Gallinago gallinago* – *G.*<br>*delicata*) | 1 | 0.50 | 1.99 | 9.45x10<sup>-10</sup> | Van Gils et al. (2020) |

**Table S3.**
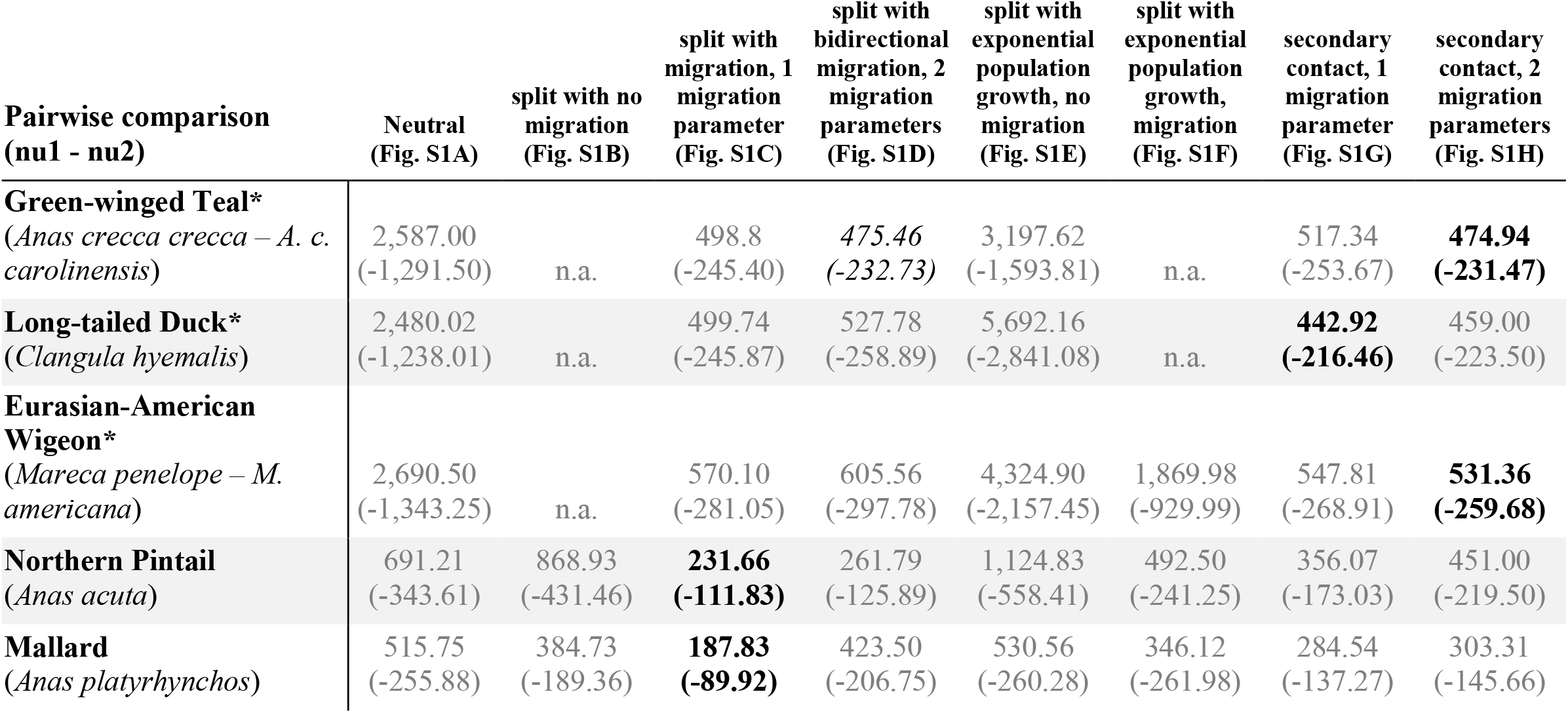
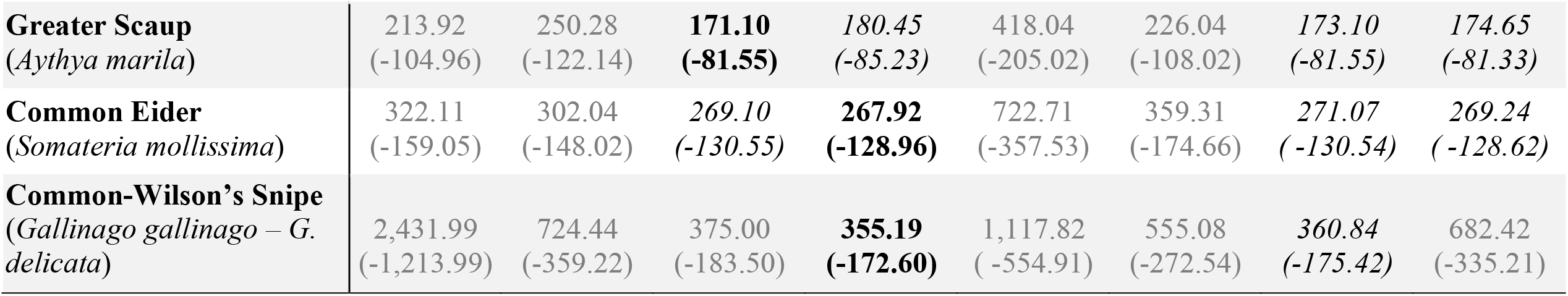
AIC and negative log-likelihood values (in parentheses) for each pairwise comparison for the eight demographic models tested (“migration” = gene flow; S1A-H refer to Supplementary Fig 1 model depictions).. When selecting best-fit models, some models had a ΔAIC of < 10, which indicates models that are not statistically separable in their likelihood of explaining the data. Calculating ΔAIC causes the best model to have ΔAIC = 0, while the rest of the models have positive values (Burnham & Anderson, 2004). Best-fit models with a weighted AIC of 1 are in bold, while runner-up models (ΔAIC < 10) are italicized. Table S5 for ΔAIC values and weighted AIC values. *Results from McLaughlin et al. (2020). Note that the “n.a.” values indicate a ΔAIC of <10. These models were unable to find a stable configuration, and thus could not be run to completion (see McLaughlin et al. 2020, Table 3).

| Pairwise comparison<br>(nu1 - nu2) | Neutral<br>(Fig. S1A) | split with no<br>migration<br>(Fig. S1B) | split with<br>migration, 1<br>migration<br>parameter<br>(Fig. S1C) | split with<br>bidirectional<br>migration, 2<br>migration<br>parameters<br>(Fig. S1D) | split with<br>exponential<br>population<br>growth, no<br>migration<br>(Fig. S1E) | split with<br>exponential<br>population<br>growth,<br>migration<br>(Fig. S1F) | secondary<br>contact, 1<br>migration<br>parameter<br>(Fig. S1G) | secondary<br>contact, 2<br>migration<br>parameters<br>(Fig. S1H) |
| --- | --- | --- | --- | --- | --- | --- | --- | --- |
| **Green-winged Teal\***<br>(*Anas crecca crecca* – *A. c. carolinensis*) | 2,587.00<br>(-1,291.50) | n.a. | 498.8<br>(-245.40) | *475.46*<br>(-232.73) | 3,197.62<br>(-1,593.81) | n.a. | 517.34<br>(-253.67) | **474.94**<br>(-231.47) |
| **Long-tailed Duck\***<br>(*Clangula hyemalis*) | 2,480.02<br>(-1,238.01) | n.a. | 499.74<br>(-245.87) | 527.78<br>(-258.89) | 5,692.16<br>(-2,841.08) | n.a. | **442.92**<br>(-216.46) | 459.00<br>(-223.50) |
| **Eurasian-American Wigeon\***<br>(*Mareca penelope* – *M. americana*) | 2,690.50<br>(-1,343.25) | n.a. | 570.10<br>(-281.05) | 605.56<br>(-297.78) | 4,324.90<br>(-2,157.45) | 1,869.98<br>(-929.99) | 547.81<br>(-268.91) | **531.36**<br>(-259.68) |
| **Northern Pintail**<br>(*Anas acuta*) | 691.21<br>(-343.61) | 868.93<br>(-431.46) | **231.66**<br>(-111.83) | 261.79<br>(-125.89) | 1,124.83<br>(-558.41) | 492.50<br>(-241.25) | 356.07<br>(-173.03) | 451.00<br>(-219.50) |
| **Mallard**<br>(*Anas platyrhynchos*) | 515.75<br>(-255.88) | 384.73<br>(-189.36) | **187.83**<br>(-89.92) | 423.50<br>(-206.75) | 530.56<br>(-260.28) | 346.12<br>(-261.98) | 284.54<br>(-137.27) | 303.31<br>(-145.66) |
| **Greater Scaup**<br>(*Aythya marila*) | 213.92<br>(-104.96) | 250.28<br>(-122.14) | **171.10**<br>**(-81.55)** | *180.45*<br>(-85.23) | 418.04<br>(-205.02) | 226.04<br>(-108.02) | *173.10*<br>(-81.55) | *174.65*<br>(-81.33) |
| **Common Eider**<br>(*Somateria mollissima*) | 322.11<br>(-159.05) | 302.04<br>(-148.02) | *269.10*<br>(-130.55) | **267.92**<br>**(-128.96)** | 722.71<br>(-357.53) | 359.31<br>(-174.66) | *271.07*<br>(-130.54) | *269.24*<br>(-128.62) |
| **Common-Wilson's Snipe**<br>(*Gallinago gallinago* – *G. delicata*) | 2,431.99<br>(-1,213.99) | 724.44<br>(-359.22) | 375.00<br>(-183.50) | **355.19**<br>**(-172.60)** | 1,117.82<br>(-554.91) | 555.08<br>(-272.54) | *360.84*<br>(-175.42) | 682.42<br>(-335.21) |

**Table S4.**
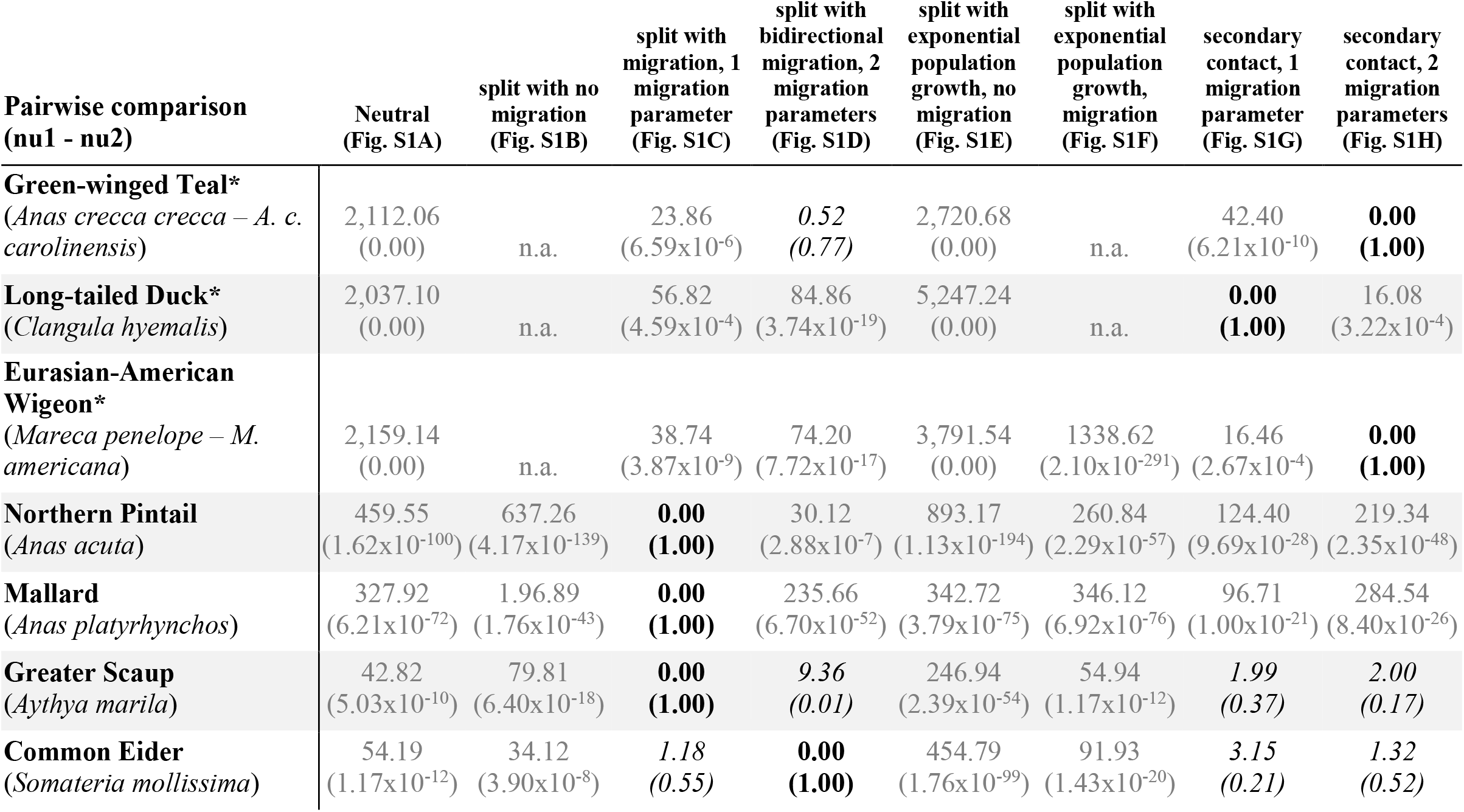
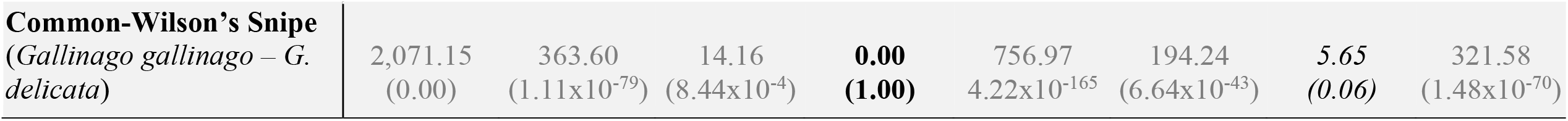
ΔAIC and weighted AIC values (in parentheses) for each pairwise comparison for the eight demographic models tested (“migration” = gene flow). Best-fit models with a ΔAIC of 0 and weighted AIC of 1 are in bold. Models that are not statistically separable from the best-fit models (ΔAIC of <10) are italicized. *Results from McLaughlin et al. (2020). “n.a.” values indicate models that were unable to find a stable configuration and thus could not be run to completion (see McLaughlin et al. 2020, Table 3).

| Pairwise comparison<br>(nu1 - nu2) | Neutral<br>(Fig. S1A) | split with no<br>migration<br>(Fig. S1B) | split with<br>migration, 1<br>migration<br>parameter<br>(Fig. S1C) | split with<br>bidirectional<br>migration, 2<br>migration<br>parameters<br>(Fig. S1D) | split with<br>exponential<br>population<br>growth, no<br>migration<br>(Fig. S1E) | split with<br>exponential<br>population<br>growth,<br>migration<br>(Fig. S1F) | secondary<br>contact, 1<br>migration<br>parameter<br>(Fig. S1G) | secondary<br>contact, 2<br>migration<br>parameters<br>(Fig. S1H) |
| --- | --- | --- | --- | --- | --- | --- | --- | --- |
| **Green-winged Teal\***<br>(*Anas crecca crecca* – *A. c. carolinensis*) | 2,112.06<br>(0.00) | n.a. | 23.86<br>(6.59x10<sup>-6</sup>) | *0.52*<br>(0.77) | 2,720.68<br>(0.00) | n.a. | 42.40<br>(6.21x10<sup>-10</sup>) | **0.00**<br>**(1.00)** |
| **Long-tailed Duck\***<br>(*Clangula hyemalis*) | 2,037.10<br>(0.00) | n.a. | 56.82<br>(4.59x10<sup>-4</sup>) | 84.86<br>(3.74x10<sup>-19</sup>) | 5,247.24<br>(0.00) | n.a. | **0.00**<br>**(1.00)** | 16.08<br>(3.22x10<sup>-4</sup>) |
| **Eurasian-American Wigeon\***<br>(*Mareca penelope* – *M. americana*) | 2,159.14<br>(0.00) | n.a. | 38.74<br>(3.87x10<sup>-9</sup>) | 74.20<br>(7.72x10<sup>-17</sup>) | 3,791.54<br>(0.00) | 1338.62<br>(2.10x10<sup>-291</sup>) | 16.46<br>(2.67x10<sup>-4</sup>) | **0.00**<br>**(1.00)** |
| **Northern Pintail**<br>(*Anas acuta*) | 459.55<br>(1.62x10<sup>-100</sup>) | 637.26<br>(4.17x10<sup>-139</sup>) | **0.00**<br>**(1.00)** | 30.12<br>(2.88x10<sup>-7</sup>) | 893.17<br>(1.13x10<sup>-194</sup>) | 260.84<br>(2.29x10<sup>-57</sup>) | 124.40<br>(9.69x10<sup>-28</sup>) | 219.34<br>(2.35x10<sup>-48</sup>) |
| **Mallard**<br>(*Anas platyrhynchos*) | 327.92<br>(6.21x10<sup>-72</sup>) | 1.96.89<br>(1.76x10<sup>-43</sup>) | **0.00**<br>**(1.00)** | 235.66<br>(6.70x10<sup>-52</sup>) | 342.72<br>(3.79x10<sup>-75</sup>) | 346.12<br>(6.92x10<sup>-76</sup>) | 96.71<br>(1.00x10<sup>-21</sup>) | 284.54<br>(8.40x10<sup>-26</sup>) |
| **Greater Scaup**<br>(*Aythya marila*) | 42.82<br>(5.03x10<sup>-10</sup>) | 79.81<br>(6.40x10<sup>-18</sup>) | **0.00**<br>**(1.00)** | *9.36*<br>(0.01) | 246.94<br>(2.39x10<sup>-54</sup>) | 54.94<br>(1.17x10<sup>-12</sup>) | *1.99*<br>(0.37) | *2.00*<br>(0.17) |
| **Common Eider**<br>(*Somateria mollissima*) | 54.19<br>(1.17x10<sup>-12</sup>) | 34.12<br>(3.90x10<sup>-8</sup>) | *1.18*<br>(0.55) | **0.00**<br>**(1.00)** | 454.79<br>(1.76x10<sup>-99</sup>) | 91.93<br>(1.43x10<sup>-20</sup>) | *3.15*<br>(0.21) | *1.32*<br>(0.52) |
| **Common-Wilson's Snipe**<br>(*Gallinago gallinago* – *G.*<br>*delicata*) | 2,071.15<br>(0.00) | 363.60<br>(1.11x10<sup>-79</sup>) | 14.16<br>(8.44x10<sup>-4</sup>) | **0.00**<br>**(1.00)** | 756.97<br>4.22x10<sup>-165</sup> | 194.24<br>(6.64x10<sup>-43</sup>) | **5.65**<br>**(0.06)** | 321.58<br>(1.48x10<sup>-70</sup>) |

**Table S5.** Raw demographic parameter output from δaδi used to calculate biological estimates. The average of the best three runs from the best-fit model for each pairwise comparison is given with the 95% confidence interval around that parameter (in parentheses). Instances of a hyphen indicated that the parameter is not present in the best-fit model. These values were translated into biological estimates using values from Table S3. Translated values are given in Table S7. *Results from McLaughlin et al. (2020).

| Pairwise comparison<br>(nu1 - nu2) | Best fit model via<br>$\delta a \delta i$ | nu1 | nu2 | m12 | m21 | m | T | T<sub>sc</sub> | Theta<br>($\theta$) |
| --- | --- | --- | --- | --- | --- | --- | --- | --- | --- |
| **Green-winged Teal\***<br>(*Anas crecca crecca* – *A. c. carolinensis*) | secondary contact, 2 migration parameters | 8.33<br>($\pm 0.54$) | 22.98<br>($\pm 0.57$) | 5.60<br>($\pm 0.15$) | 1.60<br>($\pm 0.20$) | - | 1.20<br>($\pm 0.06$) | 0.09<br>($\pm 0.05$) | 130.73<br>($\pm 3.66$) |
| **Long-tailed Duck\***<br>(*Clangula hyemalis*) | secondary contact, 1 migration parameter | 28.31<br>($\pm 0.49$) | 7.77<br>($\pm 0.47$) | - | - | 6.14<br>($\pm 0.11$) | 1.08<br>($\pm 0.22$) | 0.22<br>($\pm 0.04$) | 140.45<br>($\pm 4.58$) |
| **Eurasian-American Wigeon\***<br>(*Mareca penelope* – *M. americana*) | secondary contact, 2 migration parameters | 4.22<br>($\pm 0.20$) | 21.57<br>($\pm 0.31$) | 11.49<br>($\pm 0.23$) | 0.29<br>($\pm 0.05$) | - | 1.12<br>($\pm 0.11$) | 0.06<br>($\pm 0.16$) | 126.33<br>($\pm 39.62$) |
| **Northern Pintail**<br>(*Anas acuta*) | split with migration | 11.96<br>($\pm 0.59$) | 44.67<br>($\pm 1.56$) | - | - | 17.45<br>($\pm 0.02$) | 1.35<br>($\pm 0.06$) | - | 31.04<br>($\pm 1.18$) |
| **Mallard**<br>(*Anas platyrhynchos*) | split with migration | 17.73<br>($\pm 1.17$) | 26.74<br>($\pm 1.72$) | - | - | 6.19<br>($\pm 0.04$) | 2.26<br>($\pm 0.17$) | - | 12.86<br>($\pm 0.47$) |
| **Greater Scaup**<br>(*Aythya marila*) | split with migration | 2.58<br>($\pm 0.26$) | 3.73<br>($\pm 0.24$) | - | - | 5.39<br>($\pm 0.17$) | 0.78<br>($\pm 0.16$) | - | 19.39<br>($\pm 1.40$) |
| **Common Eider**<br>(*Somateria mollissima*) | split with bidirectional migration | 1.08<br>($\pm 0.05$) | 4.62<br>($\pm 0.24$) | 3.98<br>($\pm 0.28$) | 0.37<br>($\pm 0.05$) | - | 1.16<br>($\pm 0.13$) | - | 39.16<br>($\pm 2.84$) |
| **Common-Wilson's Snipe**<br>(*Gallinago gallinago* – *G. delicata*) | split with bidirectional migration | 2.57<br>($\pm 0.13$) | 16.10<br>($\pm 0.57$) | 0.001<br>($\pm 0.006$) | 0.53<br>($\pm 0.03$) | - | 1.47<br>($\pm 0.09$) | - | 165.61<br>($\pm 10.13$) |

**Table S6.**
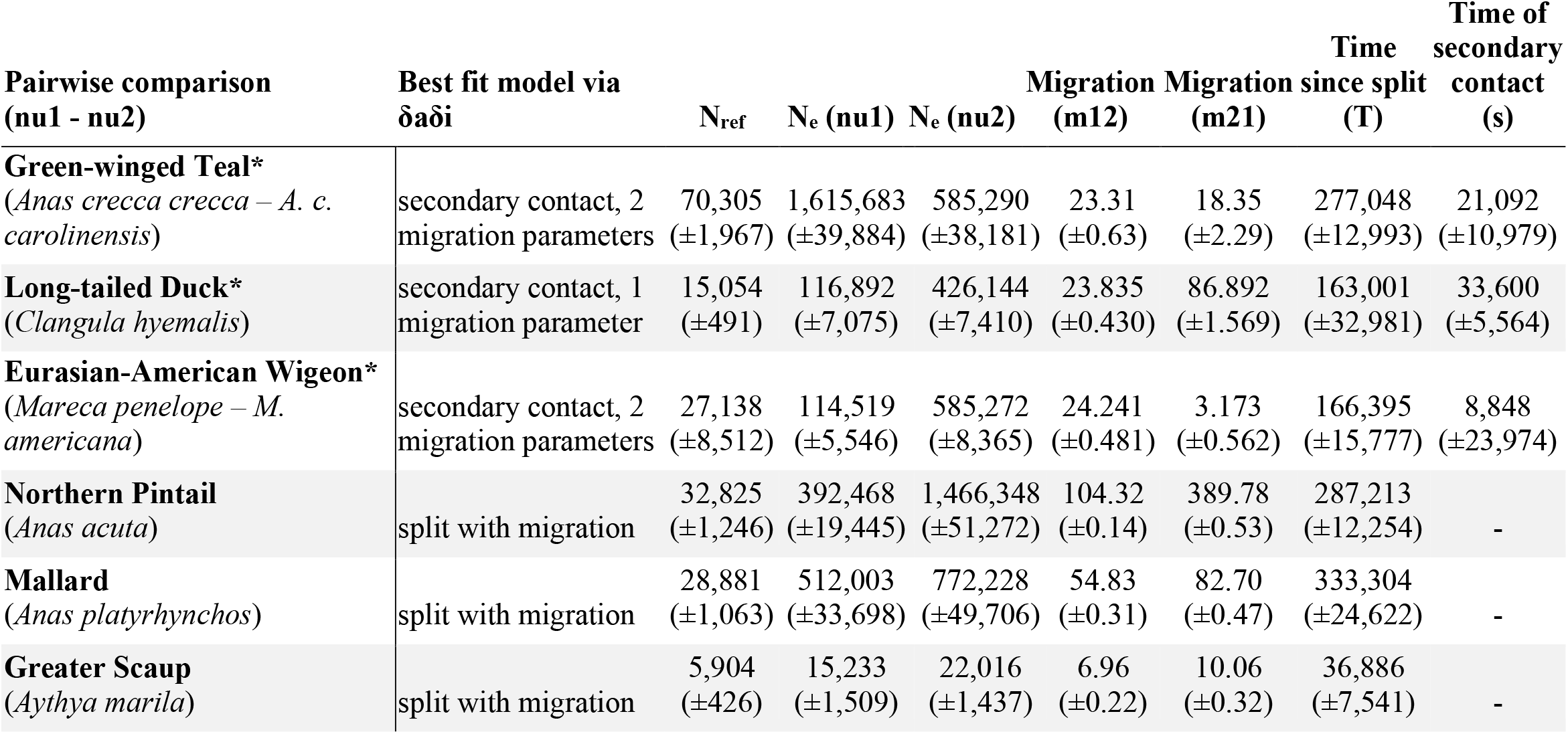
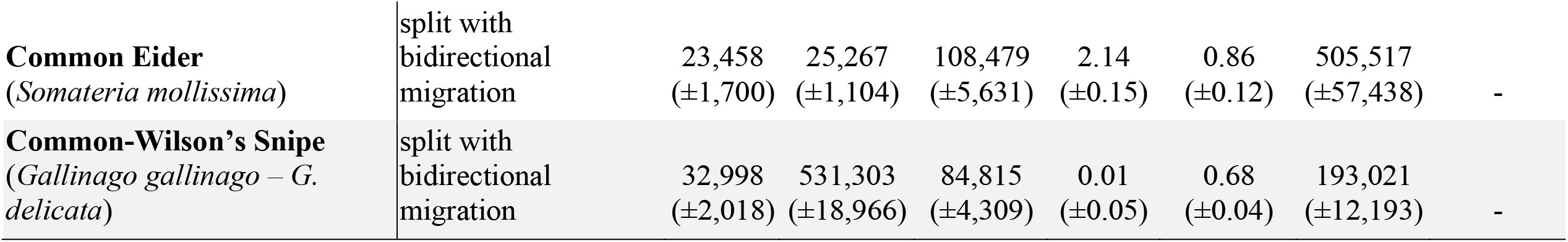
Biological estimates obtained from the best-fit δaδi models for each pairwise comparison. Here we report, as appropriate for each model, ancestral population size (*N*_*ref*_, in number of individuals), size of population 1 (*N*_*e*_ or *nu1*, in number of individuals), size of population 2 (*N*_*e*_ or *nu2*, in number of individuals), migration (gene flow) from population 1 into population 2 (*m12* as individuals/generation), migration from population 2 into population 1 (*m21*, individuals/generation), time since split (*T*, in years), and time of secondary contact (*T*_*sc*_, in years). Values in parenthesis are the ±95% confidence interval around the biological estimates. *Results from McLaughlin et al. (2020).

**Figure S1.**
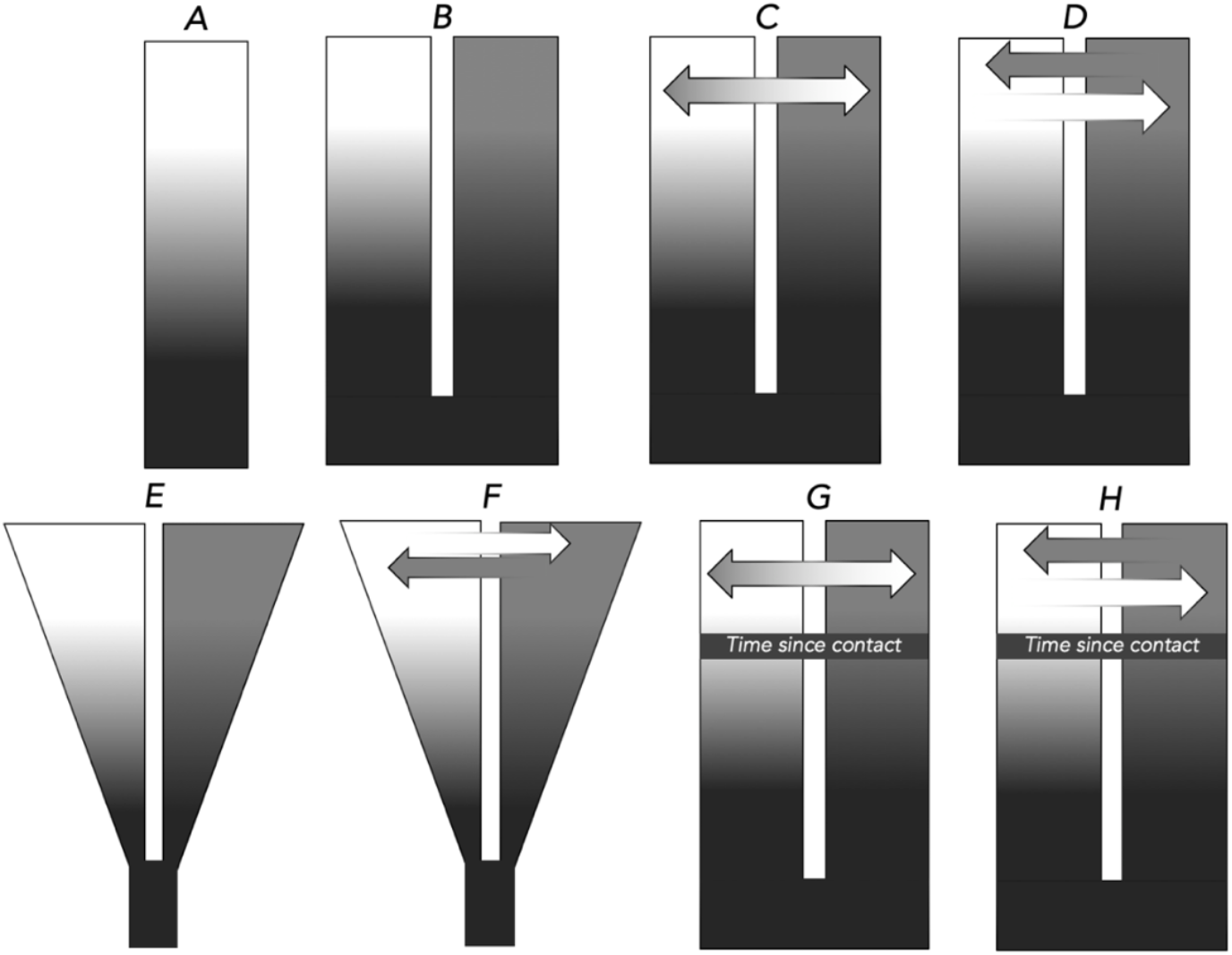
Models of divergence tested using δaδi (Gutenkunst et al., 2009) to determine demographic histories between populations: **A)** neutral, no divergence; **B)** split with no migration; **C)** split with migration, 1 migration parameter (i.e., bidirectionally symmetric); **D)** split with bidirectional migration, 2 migration parameters (i.e., bidirectionally asymmetric); **E)** split with exponential population growth, no migration; **F)** split with exponential population growth and migration; **G)** secondary contact with migration (1 migration parameter); and **H)** secondary contact with bidirectional migration (2 migration parameters). Models that contain one arrow indicate gene flow at relatively equal levels, while models with two arrows indicate unequal levels of gene flow (asymmetric). The gradient of contrast illustrates increasing population differentiation.

